# Polygenic risk score based on weight gain trajectories is predictive of childhood obesity

**DOI:** 10.1101/606277

**Authors:** Sarah J. C. Craig, Ana M. Kenney, Junli Lin, Ian M. Paul, Leann L. Birch, Jennifer S. Savage, Michele E. Marini, Francesca Chiaromonte, Matthew L. Reimherr, Kateryna D. Makova

## Abstract

Obesity is highly heritable, yet only a small fraction of its heritability has been attributed to specific genetic variants. These variants are traditionally ascertained from genome-wide association studies (GWAS), which utilize samples with tens or hundreds of thousands of individuals for whom a single summary measurement (e.g., BMI) is collected. An alternative approach is to focus on a smaller, more deeply characterized sample in conjunction with advanced statistical models that leverage detailed phenotypes. Here we use novel functional data analysis (FDA) techniques to capitalize on longitudinal growth information and construct a polygenic risk score (PRS) for obesity in children followed from birth to three years of age. This score, comprised of 24 single nucleotide polymorphisms (SNPs), is significantly higher in children with (vs. without) rapid infant weight gain—a predictor of obesity later in life. Using two independent cohorts, we show that genetic variants identified in early childhood are also informative in older children and in adults, consistent with early childhood obesity being predictive of obesity later in life. In contrast, PRSs based on SNPs identified by adult obesity GWAS are not predictive of weight gain in our cohort of children. Our research provides an example of a successful application of FDA to GWAS. We demonstrate that a deep, statistically sophisticated characterization of a longitudinal phenotype can provide increased statistical power to studies with relatively small sample sizes. This study shows how FDA approaches can be used as an alternative to the traditional GWAS.

**Author Summary:** Finding genetic variants that confer an increased risk of developing a particular disease has long been a focus of modern genetics. Genome wide association studies (GWAS) have catalogued single nucleotide polymorphisms (SNPs) associated with a variety of complex diseases in humans, including obesity, but by and large have done so using increasingly large samples-- tens or even hundreds of thousands of individuals, whose phenotypes are thus often only superficially characterized. This, in turn, may hide the intricacies of the genetic influence on disease. GWAS findings are also usually study-population dependent. We found that genetic risk scores based on SNPs from large adult obesity studies are not predictive of the propensity to gain weight in very young children. However, using a small cohort of a few hundred children deeply characterized with growth trajectories between birth and two years, and leveraging such trajectories through novel functional data analysis (FDA) techniques, we were able to produce a strong childhood obesity genetic risk score.

## Introduction

Obesity is a rising epidemic that is increasingly affecting children. In 2018, 18% of children in the United States were obese and approximately 6% were severely obese^1^—a substantial increase from previous years^2^. Given the strong association between weight gain during childhood and obesity across the life course^3, 4^, the search for early life risk factors has become a public health priority.

Obesity is a complex disease with an etiology influenced by environmental, behavioral, and genetic factors, which likely interact with each other^5^. For childhood obesity, dietary composition and sedentary lifestyle have often been cited as main contributors^6^. Evidence also exists for a significant role of parents’ socioeconomic status^7^ and maternal prenatal health factors including gestational diabetes^8^ and smoking^9^. In addition, obesity risk in children has been associated with appetite^,10^ which has been shown to be partially influenced by genetics^11^.

The heritability of obesity has been estimated to be between 50% and 90% (with the highest values reported for monozygotic twins and the lowest for non-twin siblings and parent-child pairs, reviewed in Maes *et. al*, 1997 ^12^). This is a much higher percentage than currently accounted for by known genetic variants^13, 14^. This discord is referred to as “missing heritability”—a broad discrepancy between the estimated heritability of the phenotype and the variability explained by genetic variants discovered to date. Indeed, the search for specific genetic variants that increase the risk of obesity, in adulthood as well as in childhood, is still ongoing. Using whole-genome sequencing, researchers have found variants in individual genes that contribute to severe, early-onset obesity^15^. Moreover, genome-wide association studies (GWAS) have identified single nucleotide polymorphisms (SNPs) that are significantly associated with obesity phenotypes such as increased body mass index (BMI), high waist-to-hip ratio, etc.^16–23^. Albeit successful, these studies have some shortcomings; the individual contributions of the identified SNPs tend to be very small^13^, and the prevalent focus is still on adult cohorts—with only one childhood obesity study for every 10 adult obesity studies^24^.

One way to utilize the information gained from GWAS is to summarize the risk from multiple disease-causing alleles in polygenic risk scores (PRSs) that can be computed for each individual^25^. These scores are either simple counts (unweighted) or weighted sums of disease-causing alleles identified by GWAS. Notably, while several studies have constructed PRSs for childhood obesity^23, 26–29^, most have done so relying on SNPs identified by GWAS on adult BMI. Since SNPs affecting obesity risk in adults and children may differ^30–32^, this may explain the limited^13, 23, 33^ and age-dependent^23, 34^ explanatory power of such scores for children’s weight gain status.

In this study, we contribute to bridging this gap by focusing specifically on SNPs affecting obesity risk in children. Using novel, highly effective Functional Data Analysis (FDA) techniques developed by our group on data from a small but deeply characterized pediatric cohort^35–37^, we constructed children’s growth curves and treated them as a longitudinal phenotype. FDA fully leverages this longitudinal information, extracting complex signals that can be lost in standard analyses of cross-sectional or summary measurements (e.g., BMI collected at a single time point). This increases power and specificity for assessing potentially complex and combinatorial genetic contributions. Moreover, FDA models genetic effects on the entire growth curve non-parametrically. This captures changes in effect size over time in a more flexible and effective manner than other statistical methods for longitudinal data. With our analyses, we identified genetic variants significantly associated with children’s growth curves and combined them in a novel PRS that is predictive of growth patterns and rapid infant weight gain^38, 39^, which is associated with obesity later in life. We also investigated how environmental and behavioral covariates compound with our novel score in affecting growth curves, and provided biological and statistical validations of our findings.

## Results

### Participants and DNA typing

Our study utilized 226 first-born children (out of a total of 279) enrolled in the Intervention Nurses Start Infants Growing on Healthy Trajectories (INSIGHT) study^35^. For these children, weight and length were measured at birth, 4 weeks, 16 weeks, 28 weeks, 40 weeks, and one year, and weight and height—at two and three years. Using the ratio of weight for length or height (henceforth referred to *weight-for-length/height*) at these eight time points, we constructed *growth curves* for all children (Fig. 1a; see Methods). Weight-for-length is the recommended measurement for identification of children at risk for obesity under the age of two years by the American Academy of Pediatrics (BMI is recommended afterwards)^40^. Since six out of eight time points in our study fall into this category, we utilized weight-for-length/height ratio for all eight time points analyzed for consistency.

**Figure 1.**
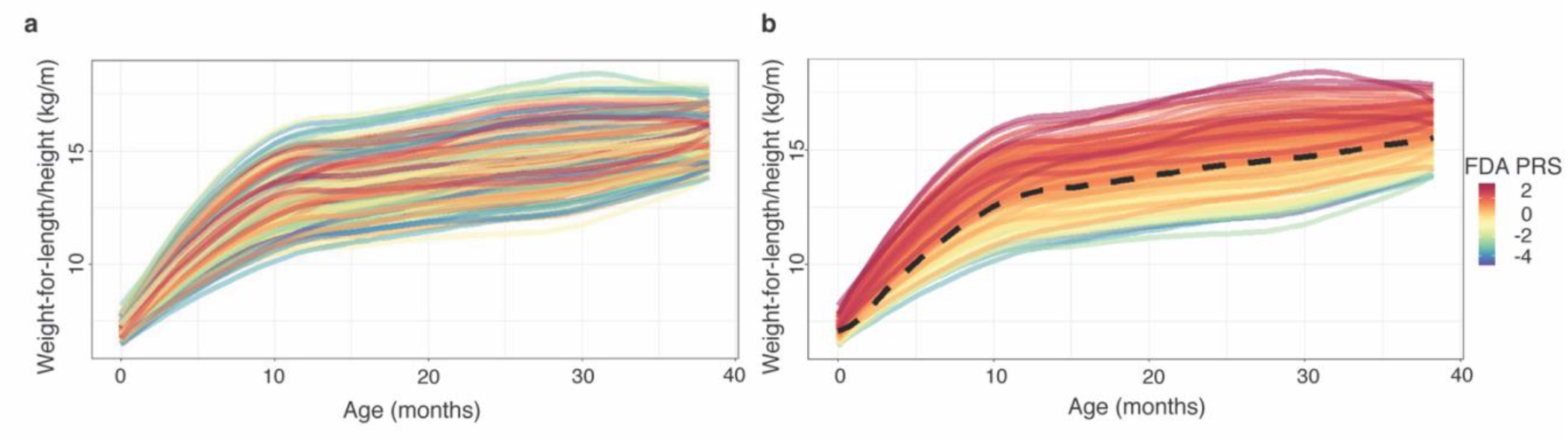
Growth curves. Growth curves from birth to three years for 226 children enrolled in the INSIGHT study are shown **(a)** color-coded by participant’s ID, and **(b)** color-coded based on a gradient corresponding to our FDA-based Polygenic Risk Score. The dashed black line is the mean curve.

In addition to growth curves, we computed the *conditional weight gain* for each child (change in weight between birth and 6 months, corrected for length, see Methods). Conditional weight gain was shown to be an effective indicator of risk for developing obesity later in life in a previous study^41^. Also in our study, children who experienced *rapid infant weight gain*, i.e. those with a positive conditional weight gain, had a significantly greater weight at one (p<2.2×10^−16^), two (p=9.1×10^−14^), and three (p=6.2×10^−13^) years of age than children who did not experience rapid infant weight gain (one-tailed t-tests, Fig. S1).

We isolated genomic DNA from blood samples from the 226 children and genotyped it on the Affymetrix Precision Medicine Research Array containing 920,744 SNPs across the genome. SNPs that had missing information, a minor allele frequency below 0.05, or were in the mitochondrial DNA were removed from the dataset—leaving a total of 329,159 SNPs for subsequent analyses (Fig. S2). With these SNPs we calculated individuals’ relatedness to assess the presence of population substructure that may confound the analysis of genomic associations. After computing a relationship matrix we regressed conditional weight gain on the top five principal components of relatedness and found no significant correlation (R^2^ =-0.003, p=0.4992). This indicates that there is no need to incorporate a population stratification into downstream analyses.

FDA-based Polygenic Risk Score predicts growth curves and rapid infant weight gain The sample size of our study is small for a traditional GWAS. However, the use of FDA techniques allowed us to leverage the longitudinal information in growth curves to identify significant SNPs and combine them into a polygenic risk score (PRS). To use FDA we restricted ourselves to a subset of 210 children and 79,498 SNP for which we had complete information (no missing values).

We first used FDA screening^42^ to reduce the analysis to the top 10,000 potentially relevant SNPs (Fig. S2). Next, we used Functional Linear Adaptive Mixed Estimation (FLAME)^43^ to identify 24 SNPs as significant predictors of children’s growth curves (Table 1). Using information from the 24 selected SNPs, we constructed our novel *FDA PRS* as a *weighted* sum of allele counts, with weights determined with additional FDA techniques (see Methods).

**Table 1.** SNPs identified as significant predictors of children’s weight gain patterns by functional data analysis. SNPs and corresponding weights in bold are the top 5 SNPs and are used in FDA5 PRS.

| SNP | Associated gene<sup>1</sup> | Chr | Position (bp) | Effect Allele | Weight | GWAS association<sup>2</sup> | Folds<sup>3</sup> | Up-down- stream gene<sup>4</sup> & associated GWAS trait<sup>2</sup> |
| --- | --- | --- | --- | --- | --- | --- | --- | --- |
| rs72679478 | *DNAJC6* | 1 | 65826847 | C | 0.111 |  | 12 | *LEPR* - early onset extreme obesity |
| rs12039940 | *ZNF648* | 1 | 181952711 | C | 0.221 |  | 17 | *CACNA1E* - longitudinal BMI measurement |
| rs10494802 | *NR5A2* | 1 | 199663441 | A | 0.152 | Waist-hip ratio | 9 |  |
| rs4915535 | *NAV1* | 1 | 201500537 | T | -0.218 | BMI, waist circumference, obesity related traits | 12 |  |
| **rs638348** | *RHOU* | 1 | 229100364 | T | **-0.358**<br>**-0.691** | Type 2 diabetes | 20 |  |
| **rs113822101** | *NOL10* | 2 | 10763339 | - | 0.230<br>**0.421** |  | 19 | *HPCAL1* - type 2 diabetes |
| rs9837708 | *FOXP1* | 3 | 71487582 | T | -0.181 | LDL cholesterol, subcutaneous adipose tissue | 14 | *EIF4E3* - eosinophil counts |
| rs921551 | *TNIP3* | 4 | 122110975 | C | -0.159 |  | 11 | *NDNF* - body weight; *ANXA5* - waist-hip ratio, type 2 diabetes; *PRDM5* - body weight |
| rs17057519 | *FABP6* | 5 | 159637815 | G | 0.115 |  | 8 | *CCNJL* - BMI-adjusted waist circumference |
| rs16889349 | *MTCH1* | 6 | 36948528 | G | 0.158 | HDL cholesterol | 18 | *FGD2* - HDL cholesterol, hypothyroidism |
| rs17626544 | *RNF144B* | 6 | 18404450 | T | 0.126 | BMI | 10 |  |
| rs4716760 | *PTPRN2* | 7 | 157376644 | T | 0.213 | Energy intake, BMI, type 2 diabetes | 19 | *UBE3C* - BMI, type 2 diabetes |
| rs1701822 | *PPP1R3A* | 7 | 112947322 | G | 0.135 | BMI | 11 |  |
| **rs62475261** | *POR* | 7 | 75533388 | C | 0.217<br>**0.396** | Eosinophil counts | 20 | *HIP1* - BMI; *RHBDD2* - eosinophil counts |
| rs10227226 | *SEC61G* | 7 | 54678150 | T | -0.322 |  | 14 | *VSTM2A* - BMI-adjusted waist-hip ratio |
| rs58307428 | *OLFM1* | 9 | 138038493 | C | 0.189 | Obesity-related traits (estradiol measurement) | 13 | *COL5A1* - waist-hip ratio |
| **rs9409226** | *BRINP1* | 9 | 122850776 | A | 0.173<br>**0.308** |  | 19 | *FBXW2* - BMI-adjusted hip circumference; *PHF19* - birth weight; *CDK5RAP2* - asthma |
| rs2389157 | *LINC00557* | 13 | 95433968 | A | 0.183 | Body composition measurement | 11 | *GPC6* - body composition measurement, visceral adipose tissue measurement |
| rs17648524 | *RBFOX1* | 16 | 7459683 | C | 0.205 | BMI, visceral adipose tissue adjusted for BMI, subcutaneous adipose tissue measurement, body weight, body composition measurement | 10 |  |
| **rs72815409** | *DNAH9* | 17 | 11751186 | A | -0.155<br>**-0.305** |  | 20 | *SHISA6* - insulin sensitivity measurement |
| rs4969367 | *BAIAP2* | 17 | 79032015 | A | 0.200 | BMI, lean body mass | 18 | *RPTOR* - BMI, obesity |
| rs141177192 | *TGIF1* | 18 | 3451388 | AA | -0.162 |  | 13 | *DLGAP1* - obesity-related traits (igfbp-3 measurement) |
| rs1539759 | *TIAM1* | 21 | 32895851 | C | 0.204 | Hypertension, renal sinus adipose tissue measurement | 17 | *SCAF4* - LDL cholesterol; *HUNK* - type 2 diabetes, periodontitis |
| rs133709 | *ELFN2* | 22 | 37827248 | A | -0.296 |  | 7 | *LGALS1* - body fat distribution |

<sup>1</sup>As determined by Affymetrix
<sup>2</sup>Association determined by the NHGRI-EBI GWAS catalog of published genome-wide association studies (https://www.ebi.ac.uk/gwas/)
<sup>3</sup>The number of times the SNP is selected in 20-fold cross validation
<sup>4</sup>Determined via LD-Link<sup>69</sup>

Our FDA PRS is indeed a strong predictor for growth curves, with a significant positive effect on weight-for-length/height ratios across time (function-on-scalar regression, in-sample R^2^=0.52, p=9.2×10^−5^; see Methods), and especially between ∼10 and ∼30 months of age (Fig. 2a). This can also be observed noting that growth curves of children with high PRS values are concentrated above the mean curve (Fig. 1b). Moreover, again in-sample, FDA PRS is significantly larger for children with rapid infant weight gain compared to those without (one-tailed t-test, p=3.3×10^−8^; Fig. 3b), and is positively correlated with conditional weight gain (R^2^=0.16, p<1×10^−05^; Fig. 3c) as well as with weight-for-length/height ratio at one (R^2^=0.50, p<1×10^−5^), two (R^2^=0.53, p<1×10^−5^), and three (R^2^=0.46, p<1×10^−5^) years of age (Fig. S3).

**Figure 2.**
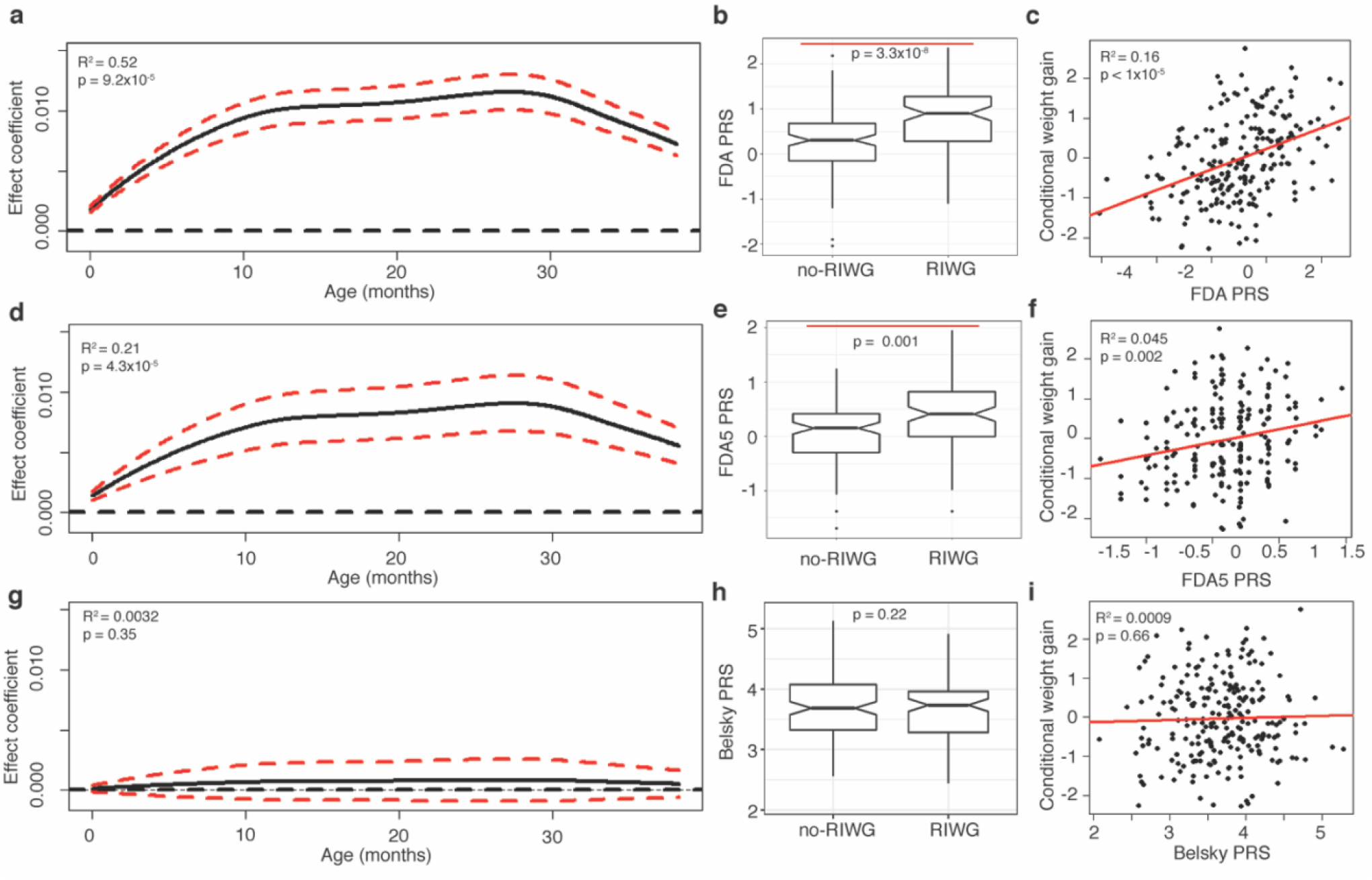
Polygenic risk scores (PRSs) and children’s growth patterns. Estimated effect coefficient for a PRS as a predictor of children’s growth curves in a function-on-scalar regression for **(a)** FDA PRS, **(d)** FDA5 PRS, or **(g)** Belsky PRS. Boxplots comparing a PRS between children with vs. without rapid infant weight gain (i.e. RIWG vs. no-RIWG) for **(b)** FDA PRS, **(e)** FDA5 PRS, or **(h)** Belsky PRS. Scatterplot of conditional weight gain vs. a PRS using **(c)** FDA PRS, **(f)** FDA5 PRS, or **(i)** Belsky PRS.

**Figure 3.**
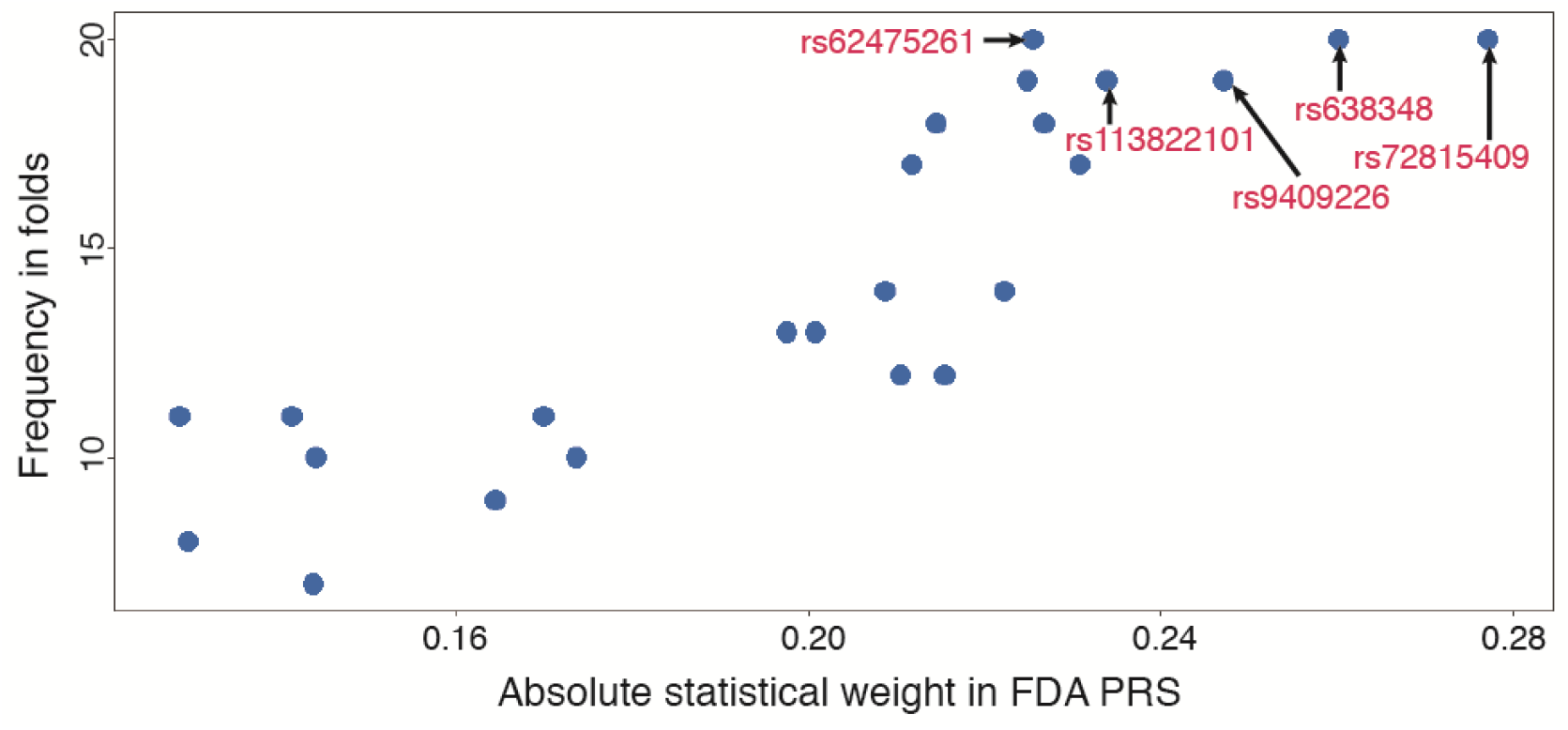
Statistical validation of FDA-based SNP selection. The frequency with which the 24 SNPs included in the FDA PRS are re-selected in a 20-fold sub-sampling scheme is plotted against their absolute statistical weight in the FDA PRS—showing a strong positive association. The SNPs with both the largest weights and the highest re-selection frequency (top five SNPs marked by arrows) may be the most important to interpret and validate in future studies.

In order to assess the robustness of our FDA-based SNP selection, we performed a sub-sampling stability analysis akin to a 20-fold cross-validation. Specifically, we randomly split the subjects into 20 equal parts (folds) and applied FLAME^43^ to perform SNP selection 20 times, with each iteration omitting a different fold. We next counted how many times (out of 20) each SNP was selected, to ascertain the stability of SNP selection. Notably, for the 24 SNPs included in our FDA PRS, the weights computed to construct the PRS correlate with the number of times the SNPs are selected in this sub-sampling scheme (Fig. 3). The frequency of selection captures how stable the effect of a genetic variant is amid the complex and combinatorial signals in this type of data. Moreover, SNPs which have both the highest selection frequency and the largest weights may be the most important to interpret and validate in future studies.

In addition to calculating a PRS based on the full complement of 24 SNPs selected by FLAME^43^, we computed a PRS restricted to the top 5 SNPs in terms of selection frequency and weight magnitude, as highlighted by our stability analysis in the previous paragraph (Fig. 3; see also Table 1). These 5 SNPs are rs72815409, rs638348, rs9409226, rs113822101, and rs62475261, and we refer to the PRS calculated on them as *FDA5 PRS*. FDA5 PRS too has a significant positive effect on weight-for-length/height ratios across time (function-on-scalar regression, in-sample R^2^=0.21, p=4.3×10^−5^; Fig. 2d), a positive correlation with conditional weight gain (R^2^ =0.045, p=0.002; Fig. 2f), and values that are significantly higher for children with rapid infant weight gain compared to those without (one-tailed t-test, p=0.001; Fig. 2e). We note here that explanatory power evaluated after model selection (in our case, post selection of SNPs) and in-sample can be highly inflated, just as in any GWAS. However, as part of the biological validation analysis presented in the section “BMI in two independent cohorts”, we confirmed the predictive performance of our scores on two completely independent data sets.

Based on the NHGRI-EBI GWAS catalog (https://www.ebi.ac.uk/gwas/), one of the FDA5 PRS SNPs is located in a gene linked to a metabolic disorder: rs638348 is located within gene *RHOU* associated with type 2 diabetes (Table 1). Additionally, two other SNPs of the FDA5 PRS are downstream of genes also associated with diabetes: rs72815409 is downstream of *SHISA6* (associated with an insulin sensitivity measurement) and rs113822101 is downstream of *HPCAL1* (associated with type 2 diabetes). The fourth SNP in FDA5 PRS, rs9409226, is upstream of *FBXW2* (associated with BMI-adjusted hip circumference), *PHF19* (associated with birth weight), and *CDK5RAP2* (associated with asthma). The fifth SNP in FDA5 PRS (rs62475261) is located upstream of *HIP1* (associated with BMI) and downstream of *RHBDD2* (associated with eosinophil counts, an asthma-related trait). Thus, rs9409226 and rs62475261 are located in the vicinity of genes associated with obesity-related traits and asthma. There is unequivocal epidemiological evidence linking obesity with asthma (reviewed in Peters *et al.* 2018 ^44^), but a shared genetic underpinning has been challenging to elucidate^45^; our results suggest further investigation is warranted.

Among the 19 SNPs included in our FDA PRS but not in the FDA5 PRS (Table 1), twelve more are located within genes linked to obesity-related traits such as BMI (rs4915535, rs471670, rs17648524, rs4969367, rs17626544, rs1701822), cholesterol levels (rs9837708 and rs16889349), body composition measurement (rs2389157), waist-to-hip ratio (rs10494802), hypertension (rs1539759), and estradiol measurement (rs58307428). Seven additional SNPs are located in the vicinity of other obesity-related genes: for example, rs72679478 is located just upstream of the leptin receptor gene (*LEPR*) which has been associated with early-onset adult obesity^46^. In summary, while none of the SNPs we identified are located in the most typical and well known obesity genes (e.g., *FTO*^34, 47^ and *MC4R*^16, 47^), all of them are located either within or in the vicinity of genes linked to obesity or metabolic disorders in previous studies.

### Sample size vs. depth

While our sample size is smaller than those of most recent GWAS studies, there is much to gain from the use of the longitudinal information in growth curves. We demonstrate this through a simulation procedure that builds upon our actual data, as to guarantee realistic settings. We considered the 210 curves employed in our analysis, and re-sampled them to create simulated populations of growth curves with different sample sizes. We associated these resampled curves to a feature akin to the PRS (referred to as “pseudo-PRS”, see Methods) sampled independently from a *N*(0,0. 5^2^). The strength of this association was calibrated on our results, using the estimated effect coefficient curve of FDA PRS (see Fig. 2a) computed on the original data. We then performed function-on-scalar regression and recorded the resulting p-value. Next, to simulate a comparable scenario with a scalar, cross-sectional response, we randomly selected one time point in each curve. We then regressed this cross-sectional response on the pseudo-PRS and the individual’s age at the time point selected, and recorded the p-value of the feature. To more realistically account for the variability in less controlled studies (e.g. based on Electronic Medical Records), we also generated cross-sectional responses with larger variation/noise, adding Gaussian errors with mean 0 and variance *s*^2^. Overall we considered three cases: no additional noise (i.e. *s*^2^=0), *s*^2^=5×10^−5^ and *s*^2^=5×10^−4^. These variances were conservatively calibrated: *s*^2^=5×10^−5^ is the within-day variability estimated from a mixed effects model on the INSIGHT data (see Methods) and *s*^2^=5×10^−4^ mimics a study where measurements are considerably less accurate. Noisy responses were also regressed on the pseudo-PRS and age, and p-values recorded. The entire procedure was repeated 100 times - producing the mean p-values and standard error bands shown in Fig. 4. We observed that a small (say n=200, close to our 210) but deeply characterized sample can be just as effective as a sample more than 4 times larger where only a cross-sectional response is measured -- even when this response contains no additional noise. Adding noise akin to that in INSIGHT increases this factor to greater than 5 times, and one would need sample sizes well beyond n=1000 to obtain comparable significance at higher levels of noise.

**Figure 4.**
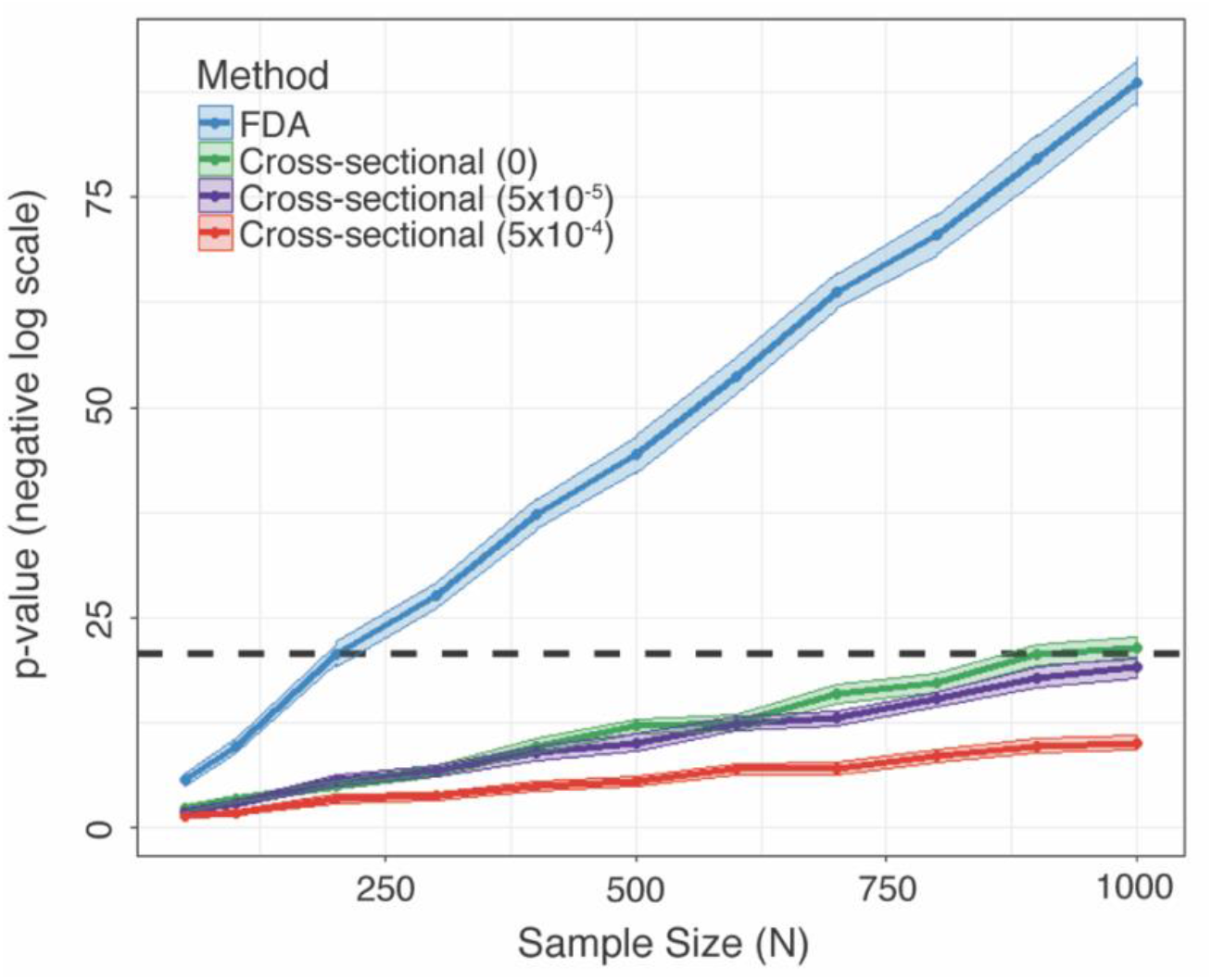
P-value curves of simulations across sample sizes. Average significance (-log(p-value)) and standard error bands as a function of the sample size, from 100 replications of a simulation procedure. Synthetic growth curves, as well as cross-sectional responses with varying levels or noise, are regressed on an artificial feature representing a polygenic risk score. The dashed black horizontal line corresponds to a p-value of 7×10^−9^, which is the one obtained using growth curves and a sample size of 200 (similar to that of our study).

### Biological Validation of the FDA-based Polygenic Risk Score

#### BMI in our cohort

To provide an initial biological validation of the FDA PRS constructed using weight-for-length/height ratio growth curves, we considered growth curves for the children in our INSIGHT cohort constructed using a different (albeit highly related) measure of weight gain, i.e. BMI. As mentioned above, weight-for-length/height ratio is recommended for children under two years of age by the American Academy of Pediatrics^40^, however, our cohort is also observed at ages two and three, when BMI is recommended as the most meaningful measurement^40^. Notably, our weight-for-length/height FDA PRS is also a strong predictor for the BMI growth curves (R^2^=0.40, p=2.9×10^−5^, function-on-scalar regression)—suggesting a reasonable consistency between the information conveyed by the two measurements, at least up to the age of three years. FDA5 PRS is a strong predictor for BMI-based growth curves as well (R^2^ =0.18, 9.1×10^−5^).

#### BMI in two independent cohorts

Among publicly available datasets, none provides genome-wide SNP data and longitudinal weight and length or height measurements for children under the age of three. Notwithstanding the unavailability of a good match to our study design, we were able to successfully validate FDA5 PRS on two independent dbGaP cohorts consisting of older children and adults. The first dataset consists of 283 children between the ages of 8 and 9 from the Philadelphia Neurodevelopment Cohort (dbGaP study phs000607.v3.p2^48–50)^ who are identified as European Americans. The average FDA5 PRS, when individuals are grouped according to BMI deciles, exhibits an increasing trend as BMI increases (Fig. 5a). Moreover, the FDA5 PRS of children in the highest BMI decile is significantly higher than in the lowest one (p=0.041, one-tailed t-test), and there is a marginally significant, positive correlation between FDA5 PRS and BMI (R^2^ =0.011, p=0.081).

**Figure 5.**
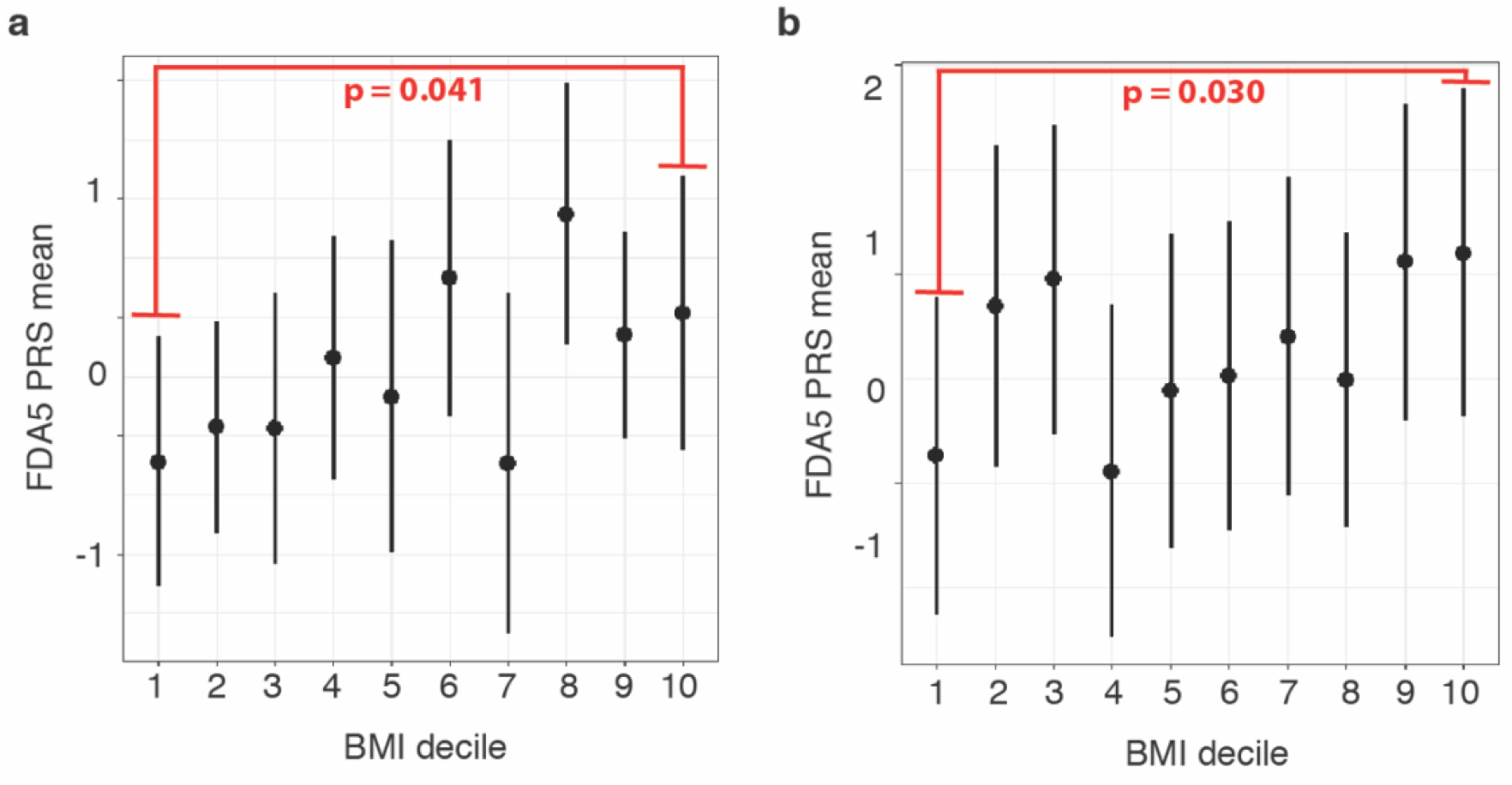
FDA-based Polygenic Risk Score and obesity in adolescent and adults validation cohorts. (**a**) Distributions of FDA5 PRS in adolescents (age 8 and 9 years) from The Philadelphia Neurodevelopment Cohort by BMI decile (n=28 per decile). (**b**) Distributions of FDA PRS in adults (over 18 years of age) either classified as normal or extremely obese in the eMERGE study by BMI decile (n=284 per decile).

The second dataset consists of 2,486 adults (≥18 years of age) from the eMERGE study (dbGaP study phs000888.v1.p1) who identify as white. We see again a significant difference in average FDA5 PRS between the lowest and highest deciles of BMI (p=0.03, one-tailed t-test; Fig. 5b). The correlation between FDA5 PRS and BMI, though still marginally significant, is weaker in this adult cohort than in the Philadelphia children’s cohort considered above (R^2^ =0.0012, p=0.087)—perhaps due to the larger difference in age with individuals in our study. Nevertheless, and remarkably, the FDA5 PRS based on our children’s weight gain patterns is predictive of extreme obesity later in life.

Considering the broader FDA PRS comprising all 24 SNPs instead of the FDR5 PRS, we did find a significant and in fact more pronounced difference between the first and 10th BMI decile in the eMERGE adults’ cohort. However, we did not find a significant difference between those BMI deciles in the Philadelphia children’s cohort (Table S1). The latter result may be due to the difficulty of validating a score based on a larger number of SNPs on small cohorts; for the 283 Philadelphia children, there are fewer allele counts across all 24 SNPs—in fact, some of the FDA PRS SNPs are completely missing. This issue does not arise for the 2,486 eMERGE adults. Notably, if we do not filter based on race and analyze the full cohort of 3,098 extremely obese and non-obese adults from eMERGE, decile differences and correlations are even more significant (Table S1). We conducted a number of other tests on these two validation cohorts (e.g., contrasting underweight and obese individuals), which further demonstrated the presence of a predictive signal in our scores (see Table S1).

### Polygenic Risk Scores based on adult obesity SNPs are not predictive of children’s growth curves and rapid infant weight gain in our cohort

While our FDA5 PRS based on children’s weight gain trajectories does validate in independent cohorts of older children and adults, PRSs based on adult GWASs do not validate in our cohort of children. First, we considered *Belsky PRS*—a weighted PRS based on 29 SNPs identified through adult obesity GWAS as described by Belsky and colleagues^28^. This PRS was shown to correlate with BMI outcomes from age three to 38, so we hypothesized it may also be a good predictor of weight outcomes in very early life. However, Belsky PRS is not a significant predictor of our children’s growth curves from birth through age three (R^2^=0.0032, p=0.35, function-on-scalar regression, Fig. 2g). Furthermore, Belsky PRS is not significantly larger for children with rapid infant weight gain compared to those without (one-tailed t-test, p=0.22; Fig. 2h) and does not display significant correlations with conditional weight gain (R^2^=0.0009, p=0.66; Fig. 2i) and weight-for-length/height ratio at one (R^2^=0.0064, p=0.25), two (R^2^=0.0036, p=0.37), and three (R^2^=0.0009, p=0.71) years of age (Fig. S4).

In addition to Belsky’s PRS, we considered four other previously published PRSs specific to childhood obesity—Elks PRS^27^, den Hoed PRS^26^, Li PRS^29^ and the recent “life-long” Khera PRS^23^ (see Methods). Similar to the Belsky PRS, the den Hoed, Li, and Khera PRSs were not significantly associated with our children’s growth curves (Fig. S5d,g, j), were not significantly different between children with vs. without rapid infant weight gain (Fig. S5e,h,k), and did not have a significant correlation with conditional weight gain (Fig. S5f,i,l). The Elks PRS showed weak but significant association with our children’s growth curves (R^2^=0.021, p=0.019; Fig. S5a) and correlation with conditional weight gain scores (R^2^=0.02, p=0.032; Fig. S5c), but no significant difference between children with vs. without conditional weight gain (two-tailed t-test, p=0.20; Fig. S5b).

### Contributions of environmental and behavioral covariates

In addition to genetics, children’s weight gain patterns can be affected by a variety of environmental and behavioral factors. To evaluate their potential effects on our results, we considered a regression of conditional weight gain^41^ on FDA PRS plus 11 potential confounding covariates, namely: maternal pre-pregnancy BMI, paternal BMI, child’s birthweight, maternal gestational weight gain, maternal gestational diabetes, maternal smoking during pregnancy, mode of delivery, the child’s gender, mother-reported child’s appetite score, INSIGHT intervention group, and family socioeconomic status (Table 2). A Best Subset Selection^51^ procedure identified only the FDA PRS (p=8.5×10^−8^) and the appetite score (p=2.4×10^−3^) as significant predictors. A regression comprising these two predictors had an R^2^ of 0.22 (Table S2), only six percentage points higher than the one obtained with FDA PRS alone (R^2^=0.16). Very similar results were obtained using FDA5 PRS in place of FDA PRS (see Table S2). However, and not unexpectedly given its lack of association with children’s growth patterns, when we reran the analysis using the Belsky PRS in place of the FDA PRS, we did not identify it as a significant predictor. Best Subset Selection for the regression of conditional weight gain on the Belsky PRS plus the 11 environmental and behavioral covariates retained only appetite as a positive and significant predictor (p=5.53×10^−5^); all other predictors, including the Belsky PRS itself, were eliminated.

**Table 2.**
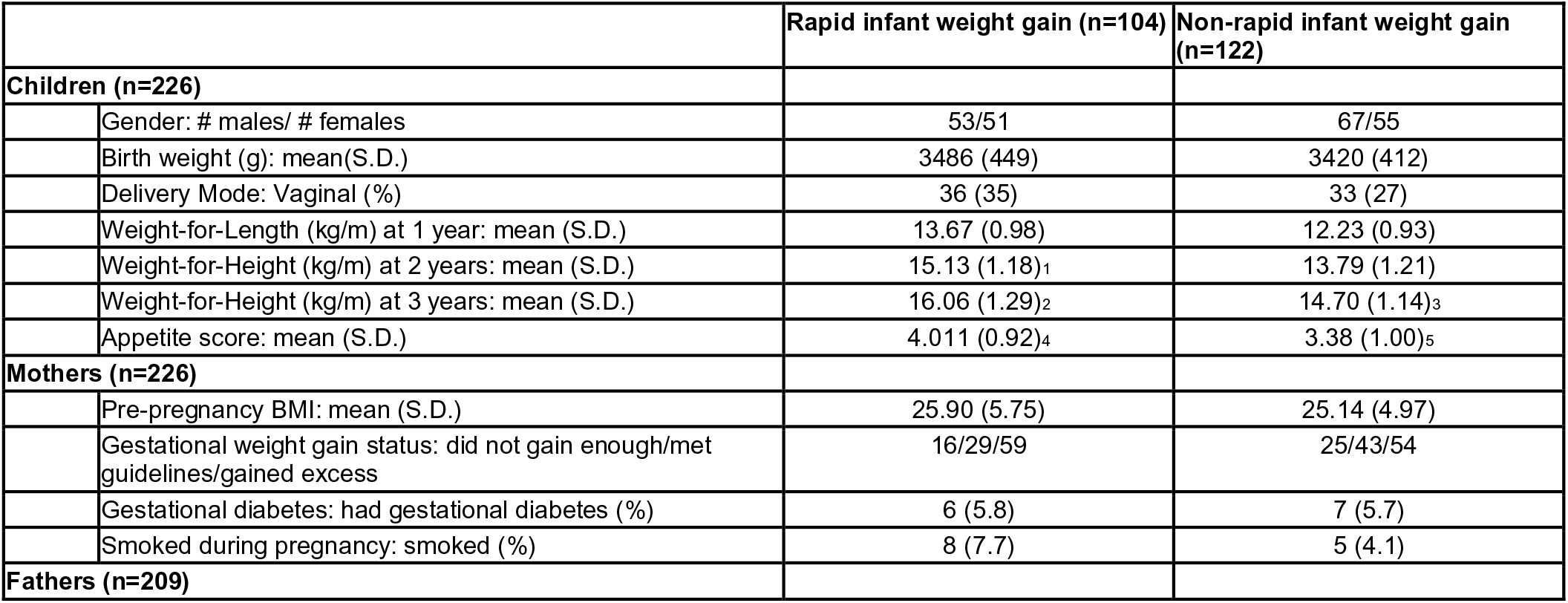
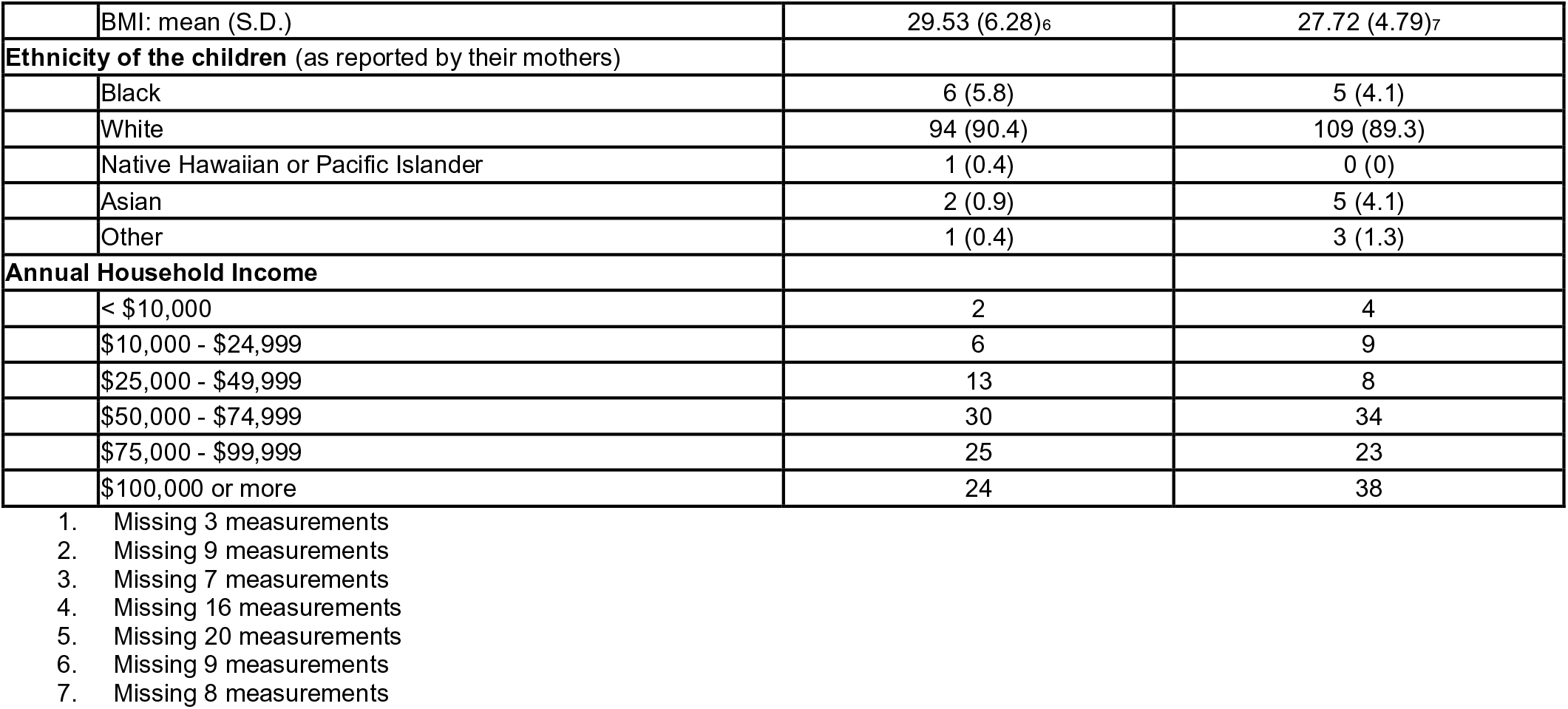
Description of the study participants.

|  | Rapid infant weight gain (n=104) | Non-rapid infant weight gain (n=122) |
| --- | --- | --- |
| **Children (n=226)** |  |  |
| Gender: # males/ # females | 53/51 | 67/55 |
| Birth weight (g): mean(S.D.) | 3486 (449) | 3420 (412) |
| Delivery Mode: Vaginal (%) | 36 (35) | 33 (27) |
| Weight-for-Length (kg/m) at 1 year: mean (S.D.) | 13.67 (0.98) | 12.23 (0.93) |
| Weight-for-Height (kg/m) at 2 years: mean (S.D.) | 15.13 (1.18)<sub>1</sub> | 13.79 (1.21) |
| Weight-for-Height (kg/m) at 3 years: mean (S.D.) | 16.06 (1.29)<sub>2</sub> | 14.70 (1.14)<sub>3</sub> |
| Appetite score: mean (S.D.) | 4.011 (0.92)<sub>4</sub> | 3.38 (1.00)<sub>5</sub> |
| **Mothers (n=226)** |  |  |
| Pre-pregnancy BMI: mean (S.D.) | 25.90 (5.75) | 25.14 (4.97) |
| Gestational weight gain status: did not gain enough/met guidelines/gained excess | 16/29/59 | 25/43/54 |
| Gestational diabetes: had gestational diabetes (%) | 6 (5.8) | 7 (5.7) |
| Smoked during pregnancy: smoked (%) | 8 (7.7) | 5 (4.1) |
| **Fathers (n=209)** |  |  |

|  |  |  |  |
| --- | --- | --- | --- |
|  | BMI: mean (S.D.) | 29.53 (6.28)<sub>6</sub> | 27.72 (4.79)<sub>7</sub> |
| **Ethnicity of the children** (as reported by their mothers) |  |  |  |
|  | Black | 6 (5.8) | 5 (4.1) |
|  | White | 94 (90.4) | 109 (89.3) |
|  | Native Hawaiian or Pacific Islander | 1 (0.4) | 0 (0) |
|  | Asian | 2 (0.9) | 5 (4.1) |
|  | Other | 1 (0.4) | 3 (1.3) |
| **Annual Household Income** |  |  |  |
| | < \$10,000 | 2 | 4 |
| | \$10,000 - \$24,999 | 6 | 9 |
| | \$25,000 - \$49,999 | 13 | 8 |
| | \$50,000 - \$74,999 | 30 | 34 |
| | \$75,000 - \$99,999 | 25 | 23 |
| | \$100,000 or more | 24 | 38 |

1. Missing 3 measurements 2. Missing 9 measurements 3. Missing 7 measurements 4. Missing 16 measurements 5. Missing 20 measurements 6. Missing 9 measurements 7. Missing 8 measurements

### Clinical translation of the FDA-based Polygenic Risk Score

A high polygenic risk score is obviously not deterministic in an individual developing a particular disease. So, it is important to assess whether an intervention on individuals with a high PRS can be successful in mitigating disease progression. Due to the nature of our children’s cohort (collected for a randomized, early-life intervention clinical trial for the prevention of obesity^35^), this question can be answered in a retrospective manner. We found that, among children with an above-average FDA PRS, those who were part of the intervention group (n=115) had a significantly lower conditional weight gain than those who were part of the control group (n= 111) (two-sided t-test, p-value = 0.036). This suggests that a screen based on the FDA PRS could potentially be used in future studies proposing intervention.

## Discussion

### Genetics of childhood and adult obesity

In this study, we used FDA techniques to construct a novel polygenic risk score (FDA PRS) which includes 24 SNPs selected based on children’s longitudinal weight gain patterns. Among our study participants, this score explains approximately 52% of the *in-sample* variability in growth curves from birth to the age of three years, and approximately 16% of the in-sample variability in conditional weight gain. We also assessed the stability of our SNP selection and constructed a second score (FDA5 PRS) using the 5 most stable SNPs among the 24. This restricted score explained approximately 21% and 4% of the in-sample variability in growth curves and conditional weight gain, respectively.

As with all genetic studies, our in-sample figures for explained variability are inflated and do not reflect predictive performance at large. We were in fact able to validate our FDA5 PRS and FDA PRS in two independent datasets comprising older children and adult individuals, but our results should still be considered preliminary. Replication in a large, prospective infant cohort would be of great benefit to show the generalizability of our risk scores as clinical markers for childhood obesity.

Interestingly, while our risk scores did validate on older individuals, none of the published PRSs based on adult BMI SNPs showed a significant association with growth curves and conditional weight gain measurements in our children cohort—with the exception of the Elks PRS ^27^, which did show a weak but detectable association signal. Notably, Elks PRS was based on SNPs identified in adult BMI GWAS but sub-selected specifically for their association with weight in children, which may explain its improved performance over other adult PRSs. Previous studies also supported only a weak relationship between PRSs based on adult BMI SNPs and childhood weight gain status^26–29^—and pointed out that the relationship became weaker the younger the age of the children^14, 26, 28^. In fact, Belsky and colleagues^28^ found no relationship between their PRS and BMI at birth and a very weak relationship at three years of age. Similarly, Khera and colleagues^23^ documented relatively weak (albeit significant) associations between their PRS and birth weight and stronger, more significant associations at age eight. These PRSs based on adult SNPs are consistent in that they show increasing effects on obesity-related phenotypes as individuals age, but almost no effect on the small children comprising our cohort. In contrast, our FDA PRSs based on childhood SNPs show a significant association with obesity-related phenotypes later in life. Even though, based on the validation datasets at our disposal, the association appears to weaken with increasing age, SNPs affecting weight gain in early childhood (i.e. those included in our scores) do retain some predictive power. This is consistent with the notion that early life weight gain, and hence its genetic underpinning, predispose to obesity across the lifecourse^3^.

The 24 SNPs identified by our study do not appear in prior PRSs for either childhood or adult obesity, except for the genome-wide Khera PRS, which contains 13 of our SNPs (as part of their set of 2,046,991 SNPs). However, all 24 are located in, or in the vicinity of, genes linked to obesity-related or metabolic disorder phenotypes in previous GWAS studies (Table 1). As with all GWAS, it is important to note that some of the identified SNPs may not be truly “causal”, but may be in linkage disequilibrium with causal SNPs—and the genes in the immediate vicinity of such SNPs may not be those through which the phenotype is influenced (e.g., rs72679478 located upstream of the leptin receptor gene). Additionally we showed that there were performance differences between the FDA PRS (with 24 SNPs) and FDA5 PRS (with only the 5 most stable SNPs from FDA PRS). Comparing the two, the FDA PRS was more effective on the growth curves and conditional weight gain score within our study sample as well as the adult validation cohort. However, FDA5 PRS successfully validated on both child and adult cohorts in spite of the smaller sample size (n = 283 for children vs 2486 for adults).

An important advantage of using a score comprising a small number of SNPs, such as ours, is that it is much more practical to compute on individuals belonging to other studies (for comparison purposes) as well as in clinical settings (for screening and potential intervention purposes). If some, potentially several, of the SNPs included in a PRS are not available for an individual, one must choose to either omit them from the calculation or identify and use proxies in their place. The larger the number of SNPs included in a score, the more SNPs may need to be omitted or proxied, reducing the fidelity of the calculation^52^ and the usefulness of the score for prediction. For instance, even having as many as 12.5 million imputed SNPs for our study cohort, we were only able to build the Khera PRS with 37% of the 2 million SNPs included in that score. In principle, this could be one explanation as to why this score did not validate in our dataset.

### The power of FDA-based GWAS

Our results demonstrate a key advantage of GWAS employing longitudinal information and FDA techniques over traditional GWAS. We faced an ultra-high dimensional problem—with many more predictors (i.e. SNPs) than observations (i.e. individuals). By integrating FDA techniques into every step of the analysis, from the screening and selection of SNPs through the construction of the PRS, we were able to utilize a more dynamic and information-rich phenotype than the ones used in traditional cross-sectional analyses. In turn, this allowed us to unveil subtler, more complex effects with a limited sample size. We illustrated this with simulations built upon our actual data -- to guarantee realistic settings. Using longitudinal data and FDA techniques, a sample size around 200 can be as effective as a cross-sectional study with more than four or five times as many individuals. This potentially expands the scope of GWAS to studies that do not comprise tens of thousands of individuals—but instead hundreds of deeply characterized participants^53^. With FDA we exploit a series of observations collected longitudinally for each individual, with benefits that include a better understanding of within-subject variability, an estimation of time-varying effects in the form of smooth curves (which reduces noise in the phenotype measurements), and a substantial gain in power. This can be very useful for investigating populations that are difficult to sample -- of course with the draw-back that collecting longitudinal data requires low participant dropout and may induce other costs or complications. Even so, this approach can provide an effective alternative to traditional GWAS, in light of the costs and benefits of collecting a single measurement on hundreds of thousands of individuals versus longitudinal measurements on a fraction of the subjects.

### Other contributing factors and perspectives

Behavioral and environmental factors are important variables to consider when investigating the etiology of complex diseases. In our study we considered 11 such factors that could influence child weight gain trajectories and found that, while the FDA PRS is by far the dominant predictor, an appetite score computed on our cohort (see Methods) has a significant effect. It has been shown that a child’s appetite behavior impacts early weight gain and may have a strong genetic basis^11, 33^. In agreement with this, a recent study found a positive relationship between a childhood obesity PRS and appetite^54^. In our study, a child’s appetite behaviour was reported by his/her mother—which could have introduced some biases. Because appetite is emerging as an interesting predictor of child weight gain status, it should be explored in more detail in future studies.

In addition to the type of environmental and behavioral factors considered in our study, other factors may interact with genetics in shaping obesity risks. These include the microbiome, the metabolome, and the epigenome. We found previously that children’s oral microbiota composition is associated with growth curves^55^. Moreover, we are collecting data on the metabolomes and epigenomes of the children in our study cohort. Our overarching goal is to develop a multi-omic model to comprehensively understand the development of childhood obesity and identify a combination of risk factors that can be used for accurate identification of children who would benefit most from early life intervention programs.

Our FDA-based polygenic risk score was computed considering the longitudinal change in weight-for-length/height ratio from birth through three years of age. An ongoing follow-up of our study participants, with weight and height collected at later time points, will allow us to further evaluate the predictive power of the FDA PRS as age progresses. Additionally, we note that our cohort (Table 2), as well as the cohorts of older children and adults used for validation, consisted predominantly of individuals of European ancestry. It will be important to conduct similar analyses on individuals of non-European ancestries, and identify differences and commonalities in the genetic factors contributing to obesity risks among different ethnicities.

The critical advantage of a PRS based on childhood vs. adult weight gain information is that the former is potentially more actionable. While INSIGHT^35^ was not designed to test an obesity intervention on individuals with high genetic risk, we were able to observe a significant weight gain velocity difference between individuals with high genetic risk in the INSIGHT intervention vs. control groups. To fully understand the clinical implications of using a PRS as a screening tool for obesity intervention additional clinical trials are needed that would combine genetic screening with early life intervention.

## Methods

### Study sample, growth curves, and conditional weight gain

We collected genetic information from 226 children recruited from the 279 families involved in the INSIGHT study^35^. These children are full-term singletons born to primiparous mothers in Central Pennsylvania. The INSIGHT study is a randomized, responsive-parenting behavioral intervention aimed at the primary prevention of childhood obesity against a home safety control. INSIGHT collected clinical, anthropometric, demographic, and behavioral variables on the children between birth and the age of three years (Table 2). In this study we utilized 11 of these variables including maternal pre-pregnancy BMI, paternal BMI, maternal pregnancy health variables (gestational weight gain, gestational diabetes, and smoking during pregnancy), family income (as a proxy for socioeconomic status), mode of delivery, child’s gender, child’s birth weight, INSIGHT intervention group (intervention or control), and mother-reported child’s appetite at 44 weeks. The appetite score is an ordinal variable on a scale from 1-5 which summarizes the Child Eating Behavior Questionnaire (CEBQ)^56^. Domains on the CEBQ include food responsiveness, emotional over-eating, food enjoyment, desire to drink, satiety responsiveness, slowness in eating, emotional under-eating, and food fussiness. Length was measured using a recumbent length board (Shorr Productions) for visits before two years (birth, 3-4 weeks, 16 weeks, 28 weeks, 40 weeks, and one year). Standing height was measured with a stadiometer (Seca 216) at two and three years.

To construct growth curves, we utilized the anthropometric data collected above to calculate weight-for-length/height ratio at each time point for our analysis. We used FDA to analyze these longitudinally as individual functions through the *fdapace* package in *R*. This package implements the Principal Analysis by Conditional Estimation (PACE) algorithm^57^, which pools information across subjects for more accurate curve construction. We used the default settings and represented them in Fig. 1a using 51 cubic spline functions with evenly spaced knots.

Conditional weight gain z-scores were calculated as the standardized residuals from a regression of age- and sex-specific weight-for-age z-score at 6-months on the weight-for-age z-score at birth (determined using the World Health Organization sex-specific child growth standards)^36^. Length-for-age z-score at 6-months, length at birth, and precise age at the 28-week visit were considered as cofactors in this regression and thus only the change in weight between birth and 6-months was captured^36, 38^. These scores are approximately normally distributed and have, by construction, a mean of 0 and a standard deviation of 1. Positive conditional weight gain z-scores correspond to a greater than average weight gain and are used to define rapid infant weight gain, which is a risk factor for developing obesity later in life^39, 58, 59^.

### Genotyping

Blood from a fingerstick was collected at the child’s one year clinical research visit. Genomic DNA was isolated (Qiagen DNeasy Blood and Tissue Kit) and genotyped on the Affymetrix Precision Medicine Research Array (PMRA). Initial quality filtering was performed using the following criteria: we removed SNPs with minor allele frequency >0.05 and/or present in less than 5% of individuals and SNPs located in mitochondrial DNA. All quality filtering steps were performed in PLINK v1.9^60, 61^ with 329,159 SNPs remaining after quality filtering.

We calculated relatedness of the INSIGHT individuals using the --make-rel command in PLINK 1.9^60, 61^. Principal components (eigenvalues and eigenvectors) of the relationship/relatedness matrix were then computed using the `eigen` function in R.

To obtain missing genotype calls and genotypes not included on the PMRA, we performed imputation. We used 294,987 of the above quality-controlled SNPs for imputation (34,172 SNPs were removed because they were not found in the 1000 Genomes Project reference). Individual’s genotypes were first phased leveraging pedigree information (genotypes were also collected for mother and father in most cases, and some younger siblings) using SHAPEIT2^62, 63^. The phased haplotypes were then used for imputation using the 1,000 Genomes Project phase 3 data^64^ as a reference panel with IMPUTE2^65^. SNPs with imputation probability <90% were removed. Following imputation, we had information for 12,479,343 SNPs.

### Functional Data Analysis techniques

First, to apply FDA techniques we further limited the number of SNPs by removing those with missing values (resulting in 210 individuals with 79,498 corresponding SNPs). After this step, we used an *FDA feature screening* method^42^, which is an effective and fast procedure to filter out SNPs that are clearly unimportant, yielding a substantially smaller subset of SNPs that can then be used in a more advanced joint model^66^. This method is specifically designed for longitudinal GWAS and can handle up to millions of SNPs. The method evaluates each SNP individually fitting a simplified model comprising only that SNP (with no other SNPs involved) and calculating a weighted mean squared error. This is then used to rank the SNPs. In our study, the top 10,000 SNPs were selected with this feature screening step. See Supplementary Note S1 for additional details about FDA feature selection.

After feature screening, we used FLAME (*Functional Linear Adaptive Mixed Estimation*)^43^, a method that simultaneously selects important predictors and produces smooth estimates for function-on-scalar linear models. This method further downselects from the pool of the top 10,000 SNPs as ranked within our screening step. In addition, it provides smooth estimates of the effects of the selected SNPs on the growth curves. To tune the penalty involved in FLAME, we split our observations into training (75%) and test (25%) sets. This procedure resulted in 24 SNPs and their corresponding estimated effect curves. See Supplementary Note S2 for additional details about FLAME.

Next, we used the estimated effect curves produced by FLAME for each of the 24 selected SNPs to construct our *FDA-based Polygenic risk score*. This was done choosing SNP-specific weights that maximize the squared covariance between weighted SNP counts and growth curves fitted through the FLAME^43^ estimates—thus incorporating both the dynamic nature of the SNP effects and linkage disequilibrium between the SNPs themselves. We applied the weights to the allele counts of each child, and computed his/her FDA PRS as the weighted sum of counts across the selected SNPs. Thus, FDA allows us to exploit the longitudinal structure of our data to not only screen and select SNPs, but also weight them using estimates of how their effects change over time.

FLAME was also used to assess the statistical robustness of SNP selection and create a more stable 5 SNP score (FDA5 PRS) through a 20-fold sub-sampling scheme where the study subjects were split at random and selection was repeated 20 times, each time on 19/20 of the data. For this exercise we fixed the penalty level in FLAME to be consistent across folds (see Supplementary Note S2 for additional details). The top 5 SNPs were identified based on their selection frequency and weight magnitude as produced by FLAME. We note that like most regularization techniques for variable selection, FLAME operates on standardized inputs (0 mean and unit variance). To generalize to other data sets, we re-scaled the weights by regressing the curves back on the raw, un-standardized values of the 24 (and 5) SNPs.

We assessed the in-sample association between growth curves and our FDA-based scores fitting function-on-scalar linear models^67^. The significance of the scores as predictors of the growth curves was determined based on three tests^68^ employing different types of weighted quadratic forms. One employs a simple L2 norm of the parameter estimate (L2), another uses principal components to reduce dimension prior to a Wald-type test (PCA), and the last blends the two through the addition of a weighted scheme in the PCA (Choi). We reported the more conservative of the three values.

### Simulation study to evaluate sample size vs depth

We created simulated populations of growth curves with different sample sizes by re-sampling the original 210 curves we used with FDA techniques (see above). In each artificial sample, we proceeded as follows. We altered the curves associating them to an artificial feature *F*, which plays the role of the PRS (the pseudo-PRS) in this abstract setting. This was done taking *Y*_*i*_(*t*) = *G*_*i*_(*t*) + *β*(*t*)*F*_*i*_ where *G*_*i*_(*t*) is the i-th original growth curve, *F*_*i*_ ∼ *N*(0,0. 5^2^) is the ith value of the artificial feature and *β*(*t*) is the estimated coefficient curve of the FDA PRS (based on 24 selected SNPs) from our original analysis. We then performed function-on-scalar regression and recorded the resulting p-value. Next, to simulate a comparable scenario with a “cross-sectional” response *Y*, we randomly selected one time point in each curve *Y*_*i*_(*t*) -- this corresponds to a measurement at a given age *t* = *A*. We then performed a standard regression of *Y* on *F* and *A*; in other words, while this regression does not use the longitudinal phenotype, it does correct for age when evaluating the effect of *F* on the cross-sectional phenotype. We recorded the p-value for *F*. Next, to simulate a larger variability occurring in less controlled or observational studies, we increased the noise adding a Gaussian error (mean 0 and variance *s*^2^) to the cross-sectional response. In a first scenario, the variance was “calibrated” on the INSIGHT data; we took *s*^2^=5×10^−5^, the within-day variation estimated from a mixed effects model for the growth index (fixed effect for age, random effects for individual and observation number -- for visits where multiple measurements were taken). In a second scenario, meant to mimic a study where measurements are less accurate than those collected in INSIGHT, we took *s*^2^= 5×10^−4^. In both scenarios we again performed a standard regression on *F* and *A*, and recorded the p-value for *F*. The whole procedure -- creating simulated samples of different sizes, generating the artificial explanatory feature (pseudo-PRS) and the various responses (growth curves), performing the various regressions and recording p-values -- was repeated 100 times, allowing us to compute averages and standard errors for the negative log p-values plotted in Fig. 4.

### Polygenic Risk Scores constructed by other studies

To calculate Belsky PRS^28^, Elks PRS^27^, den Hoed PRS^26^, Li PRS^29^, and Khera PRS^23^ on the INSIGHT cohort we employed the Allelic Scoring function in PLINK v1.9^60, 61^. For Belsky PRS, Elks PRS, and Li PRS some proxy alleles had to be used in place of SNPs that were not assessed on the PMRA. Such proxies were determined using linkage disequilibrium with LDlink^69^. Tables describing the composition of each PRS can be found in the Supplemental Materials (Tables S3-S6). We used the INSIGHT imputed data, which include 12,479,343 SNPs (see above), to calculate the Khera PRS. The Khera PRS comprises a very large number of SNPs, as many as 2,100,302^23^, but we were only able to calculate this score using 751,735 SNPs (37% of total Khera SNPs).

### Validation datasets

We used two validation datasets downloaded from dbGaP. The first dataset was obtained from Neurodevelopmental Genomics: Trajectories of Complex Phenotypes (dbGaP dataset accession number phs000607.v3.p2^48–50^). We considered 283 children between the ages of 8 and 9 years who self-reported as being of European descent. Using their height and weight measurements, we calculated BMI and then split individuals into deciles to compare those with the 10% lowest and highest BMI. BMI groups were determined using sex-specific, BMI-for-age percentiles as described by the Centers for Disease Control and Prevention. The second dataset was obtained from the eMERGE Network Imputed for 41 Phenotypes (dbGaP dataset accession number phs000888.v1.p1, variable number phv00225989.v1.p1). We considered 2,486 adults who self-identified as white. As with the children cohort we considered BMI of these individuals and categorized them based on BMI deciles (see above). For both datasets the FDA PRS was calculated using the score function in PLINK v1.9^60, 61^. Proxies for SNPs were determined using LDLink^69^ and are summarized in Table S7.

### Analysis of environmental and behavioral covariates

Using the Bayesian Information Criterion option of the `leaps` package in R^70^, we applied best subset selection^51^ to the regression of conditional weight gain scores^36^ on 11 potentially confounding covariates (described in the Results section). We included (separately) Belsky PRS, FDA PRS, or FDA5 PRS as a 12th predictor in the regression. Once the best subset of predictors was selected, we fit a linear model using the `lm` function in the R stats package.

## Data Availability

Phenotypic and Genetic data are/will be available under dbGaP study number: phs001498.v2.p1.

Code for carrying out the statistical methods (screening, applying FLAME, PRS construction and evaluation) can be found at https://github.com/makovalab-psu/InsightPRSConstruction.

## Ethics Statement

This project has been approved by Penn State IRB (PRAMS 34493). Written, informed consent was provided by mothers and/or fathers prior to a child’s enrollment.

## Acknowledgements

We are grateful for the INSIGHT study participants and nurses for their participation in this project. We would also like to thank B.Higgins, C.Reimer, R. Bruhans, A.Shelly, P.Carper, J.Beiler, J. Stokes, N.Verdiglione, and L.Hess for their assistance. <u>The Philadelphia Neurodevelopment Cohort</u>: Support for the collection of the data for Philadelphia Neurodevelopment Cohort (PNC) was provided by grant RC2MH089983 awarded to Raquel Gur and RC2MH089924 awarded to Hakon Hakonarson. Subjects were recruited and genotyped through the Center for Applied Genomics (CAG) at The Children’s Hospital in Philadelphia (CHOP). Phenotypic data collection occurred at the CAG/CHOP and at the Brain Behavior Laboratory, University of Pennsylvania. <u>eMERGE</u>: The eMERGE Network was initiated and funded by NHGRI through the following grants: U01HG006828 (Cincinnati Children’s Hospital Medical Center/Boston Children’s Hospital); U01HG006830 (Children’s Hospital of Philadelphia); U01HG006389 (Essentia Institute of Rural Health, Marshfield Clinic Research Foundation and Pennsylvania State University); U01HG006382 (Geisinger Clinic); U01HG006375 (Group Health Cooperative); U01HG006379 (Mayo Clinic); U01HG006380 (Icahn School of Medicine at Mount Sinai); U01HG006388 (Northwestern University); U01HG006378 (Vanderbilt University Medical Center); and U01HG006385 (Vanderbilt University Medical Center serving as the Coordinating Center). Samples and data in this obesity study were provided by the non-alcoholic steatohepatitis (NASH) project. Funding for the NASH project was provided by a grant from the Clinic Research Fund of Geisinger Clinic. Funding support for the genotyping of the NASH cohort was provided by a Geisinger Clinic operating funds and an award from the Clinic Research Fund. The datasets used for the analyses described in this manuscript were obtained from dbGaP at http://www.ncbi.nlm.nih.gov/gap through dbGaP accession number phs000380.v1.p1.

## Author contributions

SJCC, KDM, IMP, LLB, FC, and MR conceived the project and devised the project study design. AK, SJCC, JL, MR, FC, and KDM were involved in the data analysis. SJCC, AK, KDM, FC, and MR contributed to the writing of the manuscript with comments from co-authors. MP, LLB, JS, and MM provided resources such as access to the study population and the associated data.

