## Supplemental Information for "Polygenic risk score based on weight gain trajectories is predictive of childhood obesity"

### SUPPLEMENTARY INFORMATION:
### Polygenic risk score based on weight gain trajectories is a predictor of childhood obesity

Sarah JC Craig[1,2,#], Ana M Kenney[3,#], J Lin[3], Ian M Paul[2,4], Leann L Birch[5], Jennifer Savage[6,7], Michele E Marini[7], Francesca Chiaromonte[2,3,8]*, Matthew L Reimherr[2,3]*, Kateryna D Makova[1,2]*

1) Department of Biology, Penn State University, University Park
2) Center for Medical Genomics, Penn State University, University Park, PA
3) Department of Statistics, Penn State University, University Park, PA
4) Department of Pediatrics, Penn State College of Medicine, Hershey, PA
5) Department of Foods and Nutrition, University of Georgia, Athens, GA
6) Department of Nutritional Sciences, Penn State University, University Park, PA
7) Center for Childhood Obesity Research, Penn State University, University Park, PA
8) EMbeDS, Sant'Anna School of Advanced Studies, Piazza Martiri della Libertà, Pisa, Italy

### These authors contributed equally
*Co-corresponding authors
Kateryna Makova
Matthew Reimherr
Francesca Chiaromonte

### Content

**Content** 2

**Figure S1. Relationship between Rapid Infant Weight Gain (RIWG) status and weight-to-height ratio at one- (A), two- (B), and three-(C) years after birth.** 3

**Figure S2. Schematic of workflow, depicting the number of SNPs at each step.** 4

**Figure S3. Correlation between FDA PRS and weight-to-height ratio at one- (A), two- (B), and three- (C) years after birth.** 5

**Figure S4. Correlation between Belsky PRS and weight-to-height ratio at one- (A), two- (B), and three- (C) years after birth.** 6

**Figure S5. Relationship between INSIGHT conditional weight gain and other polygenic risk scores.** 6

**Table S1. Additional Validation Results from dbGaP datasets** 7

**Table S2. Linear regression models of FDA PRS and environmental covariates.** 9

**Table S3. Belsky PRS.** 10

**Table S4. Elks PRS.** 12

**Table S5. den Hoed PRS.** 13

**Table S6. Li PRS.** 14

**Table S7. Proxy SNPs for the FDA PRS used in calculating the score for the validation cohorts.** 17

**Note S1. FDA Feature Screening** 19

**Note S2. FLAME (Functional Linear Adaptive Mixed Estimation)** 20

**REFERENCES** 21

**Figure S1. Relationship between Rapid Infant Weight Gain (RIWG) status and weight-to-height ratio at one- (A), two- (B), and three-(C) years after birth.**

P-values determined by two-sided t-test.

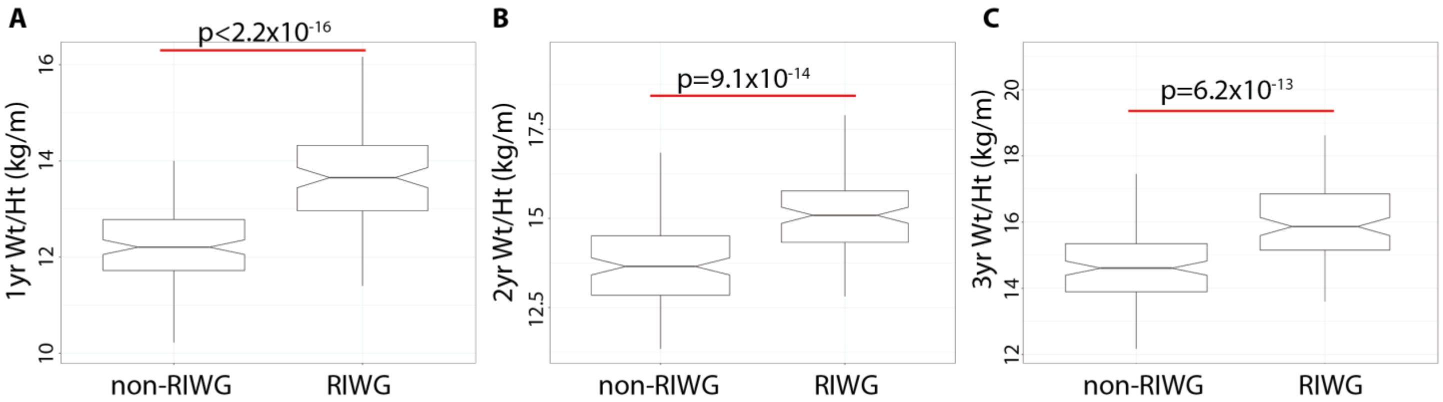

**Figure S2. Schematic of workflow, depicting the number of SNPs at each step.**

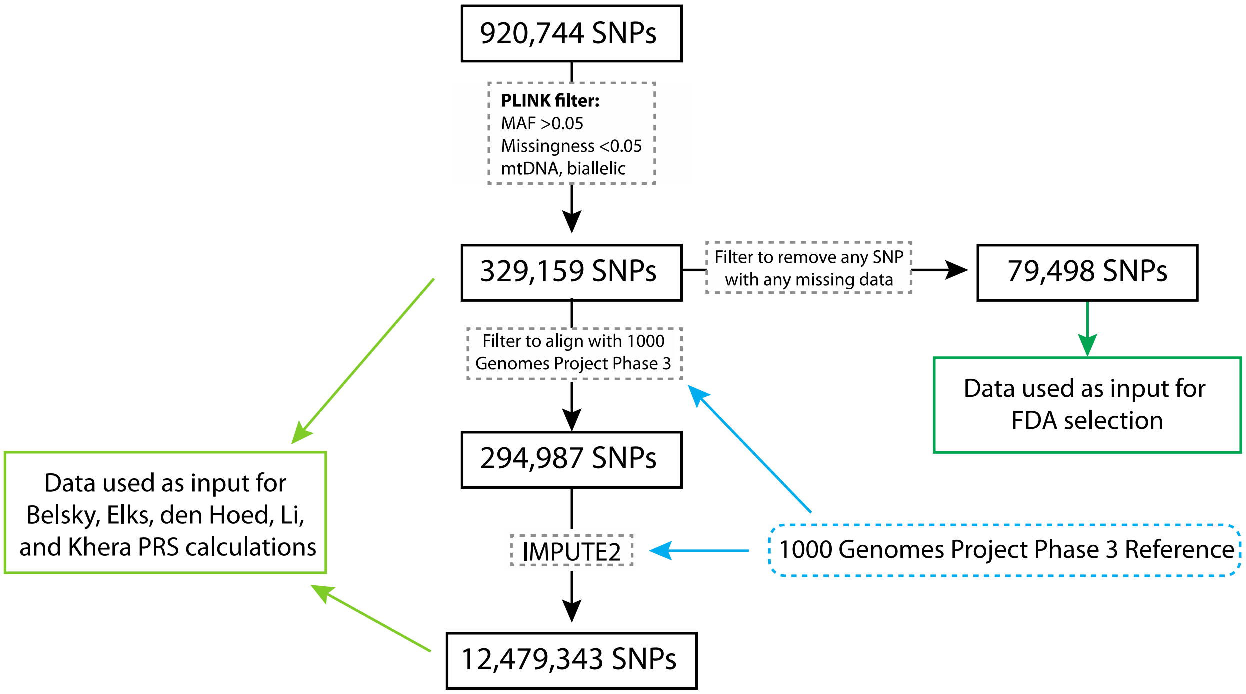

**Figure S3. Correlation between FDA PRS and weight-to-height ratio at one- (A), two- (B), and three- (C) years after birth.**

Correlations are Pearson correlations.

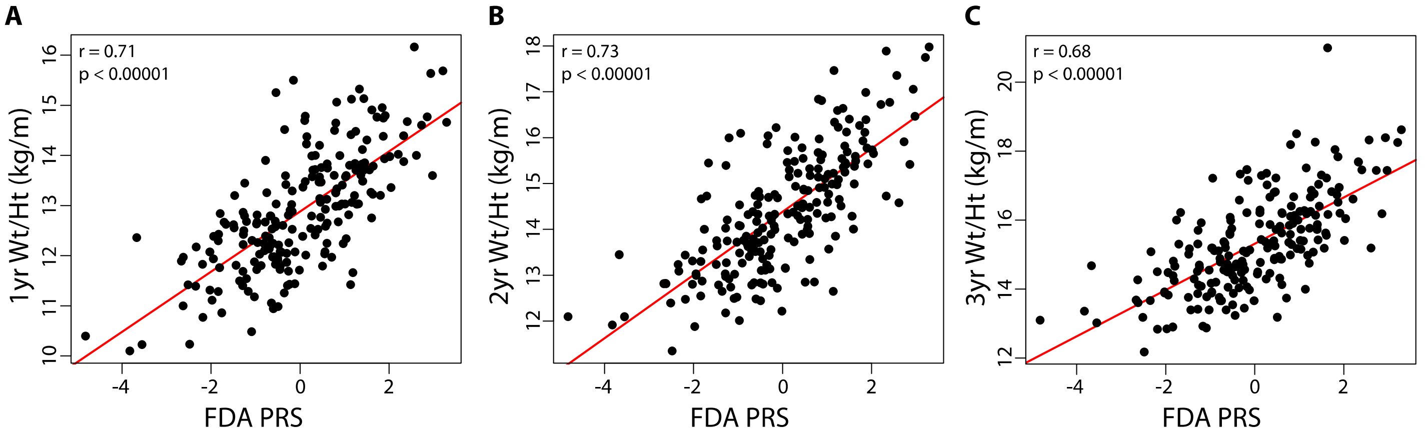

**Figure S4. Correlation between Belsky PRS and weight-to-height ratio at one- (A), two- (B), and three- (C) years after birth.**

Correlations are Pearson correlations.

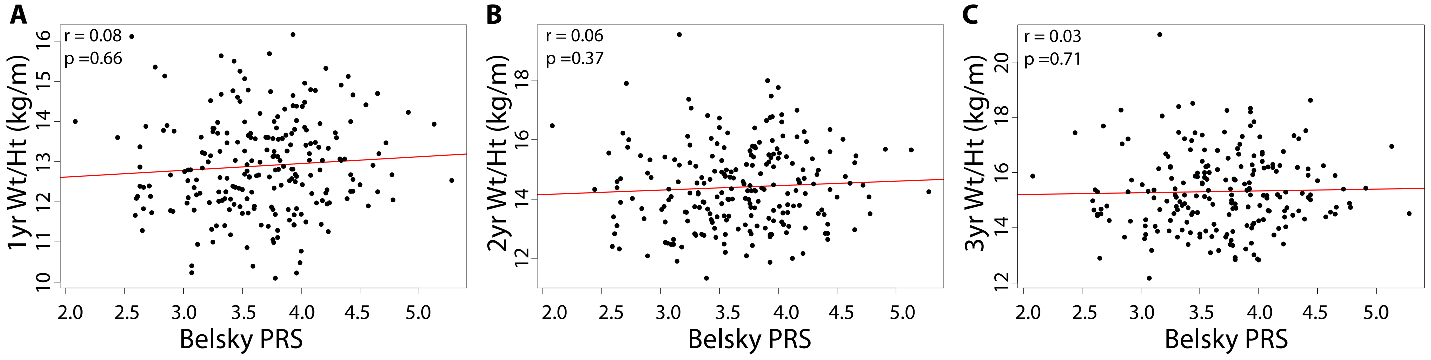

**Figure S5. Relationship between INSIGHT conditional weight gain and other polygenic risk scores.**

Function on scalar regression of growth curves and (A) Elks PRS ($R^2$ = 0.021, p = 0.019), (D) den Hoed PRS ($R^2$ = 3.5x$10^{-8}$, p = 0.95), (G) Li PRS ($R^2$ = 9.4x$10^{-9}$, p = 0.93), and (J) Khera PRS ($R^2$ = 3.54x$10^{-5}$, p = 0.79). Comparison of the (B) Elks PRS (two-sided t-test, p = 0.20), (E) den Hoed PRS (two-sided t-test, p = 0.95), (H) Li PRS (two-sided t-test, p = 0.93), and (K) Khera PRS (two-sided t-test, p = 0.64) in individuals with versus without Rapid Infant Weight Gain (RIWG). Correlation plots between (C) Elks PRS ($R^2$ = 0.019, p = 0.039), (F) den Hoed PRS ($R^2$ = 2.7x$10^{-5}$, p = 0.94), (I) Li PRS ($R^2$ = 1.2x$10^{-6}$, p = 0.99), and (L) Khera PRS ($R^2$ = 4.2x$10^{-4}$, p = 0.76) (Pearson correlations).

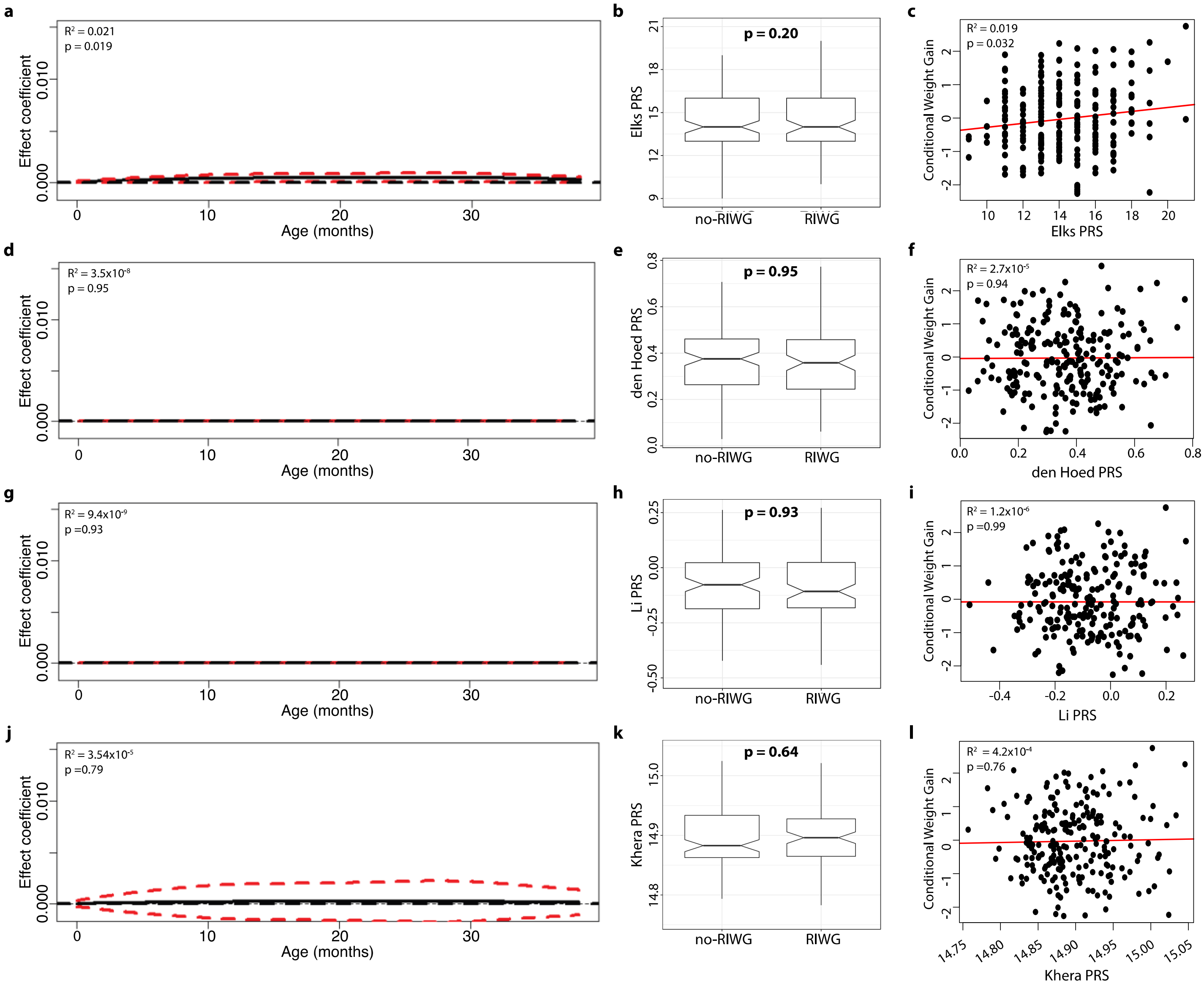

##### Table S1. Additional Validation Results from dbGaP datasets

Results from additional statistical tests performed using the Philadelphia Neurodevelopment cohort (i.e. Child cohort) and the eMERGE cohort (i.e. Adult cohort).

| | $R^2$ (p-value) regressing BMI on PRS | P-value from one-sided t-test: Compare PRS among BMI deciles | P-value from one-sided t-test: Compare BMI among PRS deciles | P-value from ANOVA: compare all BMI groups | P-value from one-sided t-test: compare extreme BMI groups |
|---|---|---|---|---|---|
| **Child cohort (n = 283)** | | **2 groups: Lowest and highest deciles** | **2 groups: Lowest and highest deciles** | **4 groups: Underweight Normal Overweight Obese** | **2 groups: Underweight Obese** |
| FDA PRS | 0.0012 (0.56) | 0.285 | 0.299 | 0.789 | 0.576 |
| FDA5 PRS | 0.011 (0.08) | 0.041 | 0.066 | 0.0857 | 0.0096 |
| **Adult cohort - white identifying (n = 2486)** | | **2 groups: Lowest and highest deciles** | **2 groups: Lowest and highest deciles** | **2 groups: Non-Obese Extremely Obese** | **N/A** |
| FDA PRS | 0.0012 (0.09) | 0.015 | 0.049 | 0.103 | |
| FDA5 PRS | 0.0012 (0.09) | 0.030 | 0.062 | 0.078 | |
| **Adult cohort - all (n = 3048)** | | **2 groups: Lowest and highest deciles** | **2 groups: Lowest and highest deciles** | **2 groups: Non-Obese Extremely Obese** | **N/A** |
| FDA PRS | 0.0027 (0.004) | 0.011 | 0.002 | 0.005 | |
| FDA5 PRS | 0.0016 (0.025) | 0.048 | 0.048 | 0.026 | |

**Table S2**. **Linear regression models of FDA PRS and environmental covariates.**

Result of linear regression models using FDA PRS, FDA5 PRS, and 11 other covariates as predictors on conditional weight gain scores as responses.

| | **Coefficient** | **p-value** | **Adjusted R-squared** |
|---|---|---|---|
| **Best model (according to best subset selection) considering FDA PRS as the genetic component** | | | |
| Appetite Score | 0.21 | 0.002 | 0.05 |
| Birthweight | -0.0002 | 0.27 | 0.007 |
| FDA PRS | 0.49 | $8.5x10^{-8}$ | 0.16 |
| Total model | | $1.9x10^{-9}$ | 0.22 |
| **Best model (according to best subset selection) considering FDA5 PRS as the genetic component** | | | |
| Appetite Score | 0.25 | 0.0006 | 0.07 |
| FDA5 PRS | 0.29 | 0.01 | 0.04 |
| Total model | | $2.5x10^{--5}$ | 0.11 |
| **Model considering only appetite score** | | | |
| Appetite Score | 0.29 | $1.3x10^{-4}$ | |
| Total model | | $1.3x10^{-4}$ | 0.084 |

**Table S3. Belsky PRS**.

Recreated from Belsky et al. 2012[1] with additional information on proxy alleles used for our population.

| Chr | Nearest Gene | rsID | Alleles | Risk Allele | GWAS Effect Size for BMI | Proxy Allele |
|---|---|---|---|---|---|---|
| 1 | *NEGR1* | rs2568958 | A/G | A | 0.13 | |
| | *TNNI3K* | rs1514177 | C/G | G | 0.07 | |
| | *PTBP2* | rs11165643 | C/T | T | 0.06 | |
| | *SEC16B* | rs10913469 | C/T | C | 0.21 | |
| 2 | *TMEM18* | rs7567570 | C/T | C | 0.31 | rs2903492: G/A (T=G, C=A), MAF=0.17, r2=0.986, D'=0.993, distance=9538 |
| | *ADCY3, RBJ* | rs101182181 | A/G | G | 0.14 | |
| | *FANCL* | rs887912 | A/G | A | 0.1 | |
| 3 | *CADM2* | rs7640855 | A/G | A | 0.1 | rs13078807: A/G (G=A, A=G), MAF=0.2087, r2=1.0, D'=1.0, distance=9986 |
| | *ETV5* | rs7647305 | C/T | C | 0.12 | |
| 4 | *SLC39A8* | rs13107325 | C/T | T | 0.19 | |
| 5 | *FLJ35779* | rs2112347 | G/T | T | 0.1 | |
| | *ZNF608* | rs6864049 | A/G | G | 0.07 | |
| 6 | *TFAP2B* | rs2206277 | A/G | A | 0.13 | |
| 9 | *LRRN6C* | rs1412235 | C/G | C | 0.11 | rs10968576: A/G (G=A, C=G), MAF=0.3022, distance=3343, D'=1.0, r2=0.986 |
| | *LMX1B* | rs867559 | A/G | G | 0.24 | |
| 11 | *STK33, RPL27A* | rs4929949 | C/T | C | 0.06 | |
| | *BDNF* | rs6265 | A/G | G | 0.18 | |
| | *MTCH2* | rs10838738 | A/G | G | 0.05 | |
| 12 | *BCDIN3, FAIM2* | rs7138803 | A/G | A | 0.12 | |
| 13 | *MTIF3* | rs1475219 | A/G | C | 0.12 | rs9581852, A/G (C=G, T=A), MAF=0.1958, r2=0.7787, D'=0.9543, distance=23667 |
| 14 | *PRKD1* | rs11847697 | C/T | T | 0.17 | |
| | *NRXN3* | rs10150332 | C/T | C | 0.13 | |

| | | | | | | |
|---|---|---|---|---|---|---|
| 15 | *MAP2K5* | rs2241423 | A/G | G | 0.13 | |
| 16 | *GPRC5B* | rs12446554 | G/T | G | 0.17 | |
| | *SH2B1* | rs4788102 | A/G | A | 0.15 | |
| | *FTO* | rs9939609 | A/T | A | 0.38 | |
| 18 | *MC4R* | rs921971 | C/T | C | 0.21 | rs8089364, T/C (T=T, C=C), MAF=0.2684, r2=1.0, D'=1.0, distance=2834 |
| 19 | *KCTD15* | rs29941 | C/T | C | 0.06 | |
| | *ZC3H4, TMEM160* | rs3819291 | A/G | A | 0.09 | |

**Table S4. Elks PRS.**

Recreated from Elks *et al.*[2] with additional information on proxy alleles used for our population. No effect size is reported because this is an unweighted PRS. The score is calculated as the sum of the number of effect alleles.

| SNP | Nearest Gene | Effect Allele | Proxy used |
|---|---|---|---|
| rs10146997 | *NRXN3* | *G* | |
| rs13107325 | *SLC39A8* | *T* | |
| rs1514175 | *TNNI3K* | *A* | |
| rs1555543 | *PTBP2* | *C* | |
| rs17782313 | *MC4R* | *C* | |
| rs2112347 | *FLJ35779* | *T* | |
| rs2568958 | *NEGR1* | *A* | |
| rs4836133 -- chr 5 | *ZNF608* | *A* | |
| rs4929949 | *RPL27A* | *C* | |
| rs6548238 | *TMEM18* | *C* | |
| rs713586 | *RBJ/POMC* | *C* | |
| rs7640855 | *CADM2* | *A* | rs13078807, A/G (G=A,A=G),MAF = 0.2087, $r^2$ = 1.0, D' = 1.0, distance = 9986 |
| rs7647305 | *TRA2B* | *C* | |
| rs925946 | *BDNF* | *T* | |
| rs987237 | *TFAP2B* | *G* | |
| rs9941349 | *FTO* | *T* | |

**Table S5. den Hoed PRS.**

Recreated from den Hoed *et al.*[3] with additional information on proxy alleles used for our population.

| SNP | Nearest Gene | Effect Size | Effect Allele | Proxy used |
|---|---|---|---|---|
| rs2815752 | *NEGR1* | 0.081 | G | |
| rs10913469 | *SEC16B* | 0.136 | C | |
| rs2605100* | *LYPLAL1* | 0.012 | A | |
| rs6548238 | *TMEM18* | 0.148 | T | |
| rs7647305 | *ETV5* | 0.048 | T | |
| rs10938397 | *GNPDA2* | 0.051 | G | |
| rs987237 | *TFAP2B* | 0.069 | G | |
| rs545854 | MSRA | -0.08 | G | |
| rs1488830 | BDNF | 0.037 | C | rs4074134, C/T (C=T,T=C), MAF = 0.2187, $r^2$ = 0.9942, D'= 1.0, distance = 10400 |
| rs925946 | BDNF | 0.057 | T | |
| rs10838738 | MTCH2 | -0.017 | G | |
| rs7138803 | *BCDIN3D* | 0.045 | A | |
| rs10146997 | *NRXN3* | 0.022 | G | |
| rs8055138 | *SH2B1* | 0.012 | T | |
| rs1121980 | *FTO* | 0.02 | A | |
| rs17782313 | *MC4R* | 0.013 | C | |
| rs11084753 | *KCTD15* | 0.02 | A | |

**Table S6. Li PRS.**

Recreated from Li *et al.*[4] with additional information on proxy alleles used for our population.

| SNP | Nearest Gene | Effect Size | Effect Allele | Proxy used |
|---|---|---|---|---|
| rs1000940 | *RABEP1* | 0.0192 | G | |
| rs10132280 | *STXBP6* | -0.023 | C | rs8015400, C/A (C=A, A=C), MAF=0.3479, $r^2$=0.8904, D'=1.0, distance=2809 |
| rs10150332 | *NRXN3* | 0.024 | A | |
| rs10733682 | *LMX1B* | 0.0174 | A | |
| rs10838738 | *MTCH2* | -0.0241 | G | |
| rs10938397 | *GNPDA2* | -0.0402 | G | |
| rs10968576 | *LRRN6C* | 0.0249 | G | |
| rs11057405 | *CLIP1* | -0.0307 | G | |
| rs11084753 | *KCTD15* | -0.0165 | G | |
| rs11126666 | *KCNK3* | -0.0207 | A | |
| rs11191560 | *NT5C2* | -0.0308 | C | |
| rs11583200 | *ELAVL4* | 0.0177 | C | |
| rs1167827 | *HIP1* | -0.0202 | G | |
| rs11688816 | *EHBP1* | -0.0172 | G | |
| rs11727676 | *HHIP* | -0.0358 | T | |
| rs11847697 | *PRKD1* | 0.0492 | T | |
| rs12286929 | *CADM1* | 0.0217 | G | |
| rs12444979 | *GPRC5B* | -0.0396 | C | |
| rs13078807 | *CADM2* | -0.0284 | G | |
| rs13107325 | *SLC39A8* | -0.0477 | T | |
| rs13191362 | *PARK2* | 0.0277 | A | |
| rs13201877 | *IFNGR1* | -0.0233 | G | |
| rs1441264 | *MIR548A2* | 0.0175 | A | |
| rs1460676 | *FIGN* | -0.0197 | C | |
| rs1514175 | *TNNI3K* | 0.023 | A | |
| rs1528435 | *UBE2E3* | 0.0178 | T | |
| rs1555543 | *PTBP2* | -0.0238 | T | |
| rs16851483 | *RASA2* | -0.0483 | G | |
| rs16907751 | *ZBTB10* | 0.035 | C | |
| rs17001654 | *SCARB2* | -0.0306 | A | |

| | | | | |
|---|---|---|---|---|
| rs17094222 | *HIF1AN* | 0.0249 | C | |
| rs17203016 | *CREB1* | 0.021 | G | |
| rs17724992 | *PGPEP1* | 0.0194 | A | |
| rs1928295 | *TLR4* | -0.0188 | T | |
| rs2033529 | *TDRG1* | 0.019 | G | |
| rs2033732 | *RALYL* | 0.0192 | C | |
| rs206936 | *HMGA1* | -0.0186 | G | |
| rs2075650 | *TOMM40-APOE-APOC1* | 0.0258 | A | |
| rs2080454 | *CBLN1* | -0.0168 | C | |
| rs2112347 | *FLJ35779* | -0.0261 | T | |
| rs2176598 | *HSD17B12* | 0.0198 | T | |
| rs2241423 | *MAP2K5* | -0.031 | G | |
| rs2287019 | *GIPR* | 0.036 | C | |
| rs2365389 | *FHIT* | 0.02 | C | |
| rs2650492 | *SBK1* | 0.0207 | A | |
| rs2815752 | *NEGR1* | -0.0326 | A | |
| rs2820292 | *NAV1* | -0.0195 | C | |
| rs2836754 | *ETS2* | 0.0164 | C | |
| rs3736485 | *DMXL2* | 0.0176 | A | |
| rs3810291 | *TMEM160* | 0.0283 | A | |
| rs4740619 | *C9orf93* | 0.0179 | T | |
| rs4787491 | *INO80E* | -0.0159 | G | |
| rs492400 | *USP37* | -0.0158 | C | |
| rs4929949 | *TUB* | 0.0173 | C | |
| rs543874 | *SEC16B* | 0.0482 | G | |
| rs571312 | *MC4R* | 0.0553 | A | |
| rs6265 | *BDNF* | 0.0424 | G | |
| rs6465468 | *ASB4* | -0.0166 | T | rs763464, C/T G=T, T=C, MAF=0.3459, $r_2$=0.758, D'=0.9599, distance=7685 |
| rs6477694 | *EPB41L4B* | 0.0174 | C | |
| rs6548238 | *TMEM18* | -0.0605 | C | |
| rs657452 | *AGBL4* | 0.0227 | A | |
| rs6804842 | *RARB* | -0.0185 | G | |
| rs713586 | *POMC* | -0.0302 | G | |

| | | | |
|---|---|---|---|
| rs7138803 | *FAIM2* | -0.0315 | A |
| rs7164727 | *LOC100287559* | 0.018 | T |
| rs7239883 | *LOC284260* | 0.0164 | G |
| rs7243357 | *GRP* | -0.0217 | T |
| rs7498665 | *SH2B1* | 0.0307 | G |
| rs758747 | *NLRC3* | -0.0225 | T |
| rs7599312 | *ERBB4* | 0.022 | G |
| rs7647305 | *ETV5* | -0.0358 | C |
| rs7899106 | *GRID1* | -0.0395 | G |
| rs7903146 | *TCF7L2* | -0.0234 | C |
| rs887912 | *FANCL* | 0.0228 | T |
| rs925946 | *BDNF* | 0.0293 | T |
| rs9374842 | *LOC285762* | 0.0187 | T |
| rs9400239 | *FOXO3* | 0.0188 | C |
| rs9540493 | *MIR548X2* | -0.0172 | A |
| rs977747 | *TAL1* | 0.0167 | T |
| rs987237 | *TFAP2B* | 0.044 | G |
| rs9914578 | *SMG6* | 0.0201 | G |
| rs9925964 | *KAT8* | -0.0192 | A |
| rs9939609 | *FTO* | 0.0776 | A |

**Table S7. Proxy SNPs for the FDA PRS used in calculating the score for the validation cohorts.**

| SNP | Proxy for phs000607: Philadelphia Neurodevelopment Cohort | Proxy for phs000888: eMERGE |
|---|---|---|
| rs72679478 | | |
| rs12039940 | rs11803436, A/G (T=A,C=G), MAF = 0.317, r2 = 0.737, D' = 0.893, distance = 86731 | |
| rs10494802 | rs1429700, C/T (G=C,A=T), MAF = 0.192, r2 = 0.779, D' = 0.912, distance = 25095 | |
| rs4915535 | rs4915537, C/T (G=C,T=T), MAF = 0.126, r2 = 0.991, D' = 1.0, distance = 288 | |
| rs638348 | rs603404, A/G (T=A,C=G), MAF = 0.322, r2 = 0.842, D' = 0.949, distance = 4781 | |
| rs113822101 | rs715247, A/G (-=A,C=G), MAF = 0.483, r2= 0.988, D' = 1.0, distance = 27048 | rs12464817, A/G (-=A, C=G), MAF = 0.483, r2 = 0.988, D' = 1.0, distance = 56127 |
| rs9837708 | | |
| rs921551 | | |
| rs17057519 | | |
| rs16889349 | rs9470449, T/C (A=T,G=C), MAF = 0.0885, r2 = 0.918, D' = 0.988, distance = 1142 | |
| rs17626544 | | |
| rs4716760 | | |
| rs1701822 | | |
| rs62475261 | rs10231459, T/G (T=T,C=G), MAF = 0.227, r2 = 1.0, D' = 1.0, distance = 4098 | |
| rs10227226 | rs1454520, G/A (C=G,T=A), MAF = 0.428, r2 = 0.995 , D' = 1.0, distance = 2753 | |

| | | |
|---|---|---|
| rs58307428 | rs17495030, C/T (T=C,C=T), MAF = 0.271, r2 = 0.8048, D' = 1.0, distance = 400 | |
| rs9409226 | | |
| rs2389157 | | |
| rs17648524 | rs8049549, A/C (G=A,C=C), MAF = 0.387, r2 = 0.732, D' = 0.990, distance = 1840 | |
| rs72815409 | rs4792189, C/T (G=T,A=C), MAF = 0.0596, r2 = 0.756, D' = 0.9575, distance = 22574 | |
| rs4969367 | | |
| rs141177192 | | rs238534, C/T (-=C,AA=T), MAF = 0.063, r2 = 0.795, D' = 1, distance = 456 |
| rs1539759 | rs2833427, T/C (C=C,T=T), MAF = 0.361, r2 = 0.652, D' = 0.848, distance = 37430 | |
| rs133709 | | |

#### Note S1. FDA Feature Screening

We applied a feature screening method[5] to rank the SNPs based on their relevance to the phenotypes in order to quickly and efficiently remove clearly unimportant SNPs. This approach was constructed for screening ultrahigh dimensional longitudinal data, naturally fitting our setting. We considered marginal models for each SNP $j$:

$$Growth_i(t) = \beta_0(t) + \beta_{gender}(t)Gender_i + \gamma_j(t)SNP_j + \varepsilon_i(t)$$

Where the growth curve responses, $Growth_i(t)$, were generated through the weight-for-length/height ratios of each child. We computed a weighted mean squared error for each marginal model and used these to rank the SNPs (smaller error corresponding to higher ranks). We compared **weighted** mean squared error in particular since the method takes into account within subject correlation, or rather, time-varying error variance. The top 10,000 SNPs were then selected based on this criteria and used for future analysis. Overall this step brought us down from 79,498 to 10,000 SNPs under consideration for PRS construction.

#### Note S2. FLAME (Functional Linear Adaptive Mixed Estimation)

We used FLAME[6] after screening to further downselect from the pool of 10,000 SNPs to select the SNPs most informative to the phenotype. This approach can be thought of as a generalization of adaptive LASSO[7] to functional or longitudinal outcomes. FLAME simultaneously selects important features and produces smooth estimates, $\beta$, by minimizing the target:

$$\frac{1}{2N}||Y - X\beta||_{\mathbb{H}}^{2} + \lambda \sum_{i}^{I} \widetilde{w_i}||\beta_i||_{\mathbb{K}}$$

With $Y \in \mathbb{H}^N$ and $X \in \mathbb{R}^{N \times I}$ where $\mathbb{H}$ is a real separable Hilbert space with norm $||\cdot||_{\mathbb{H}}$ and $\mathbb{K}$ is a subspace of $\mathbb{H}$:

$$\mathbb{K} := \{h \in \mathbb{H}: \sum_{i=1}^{\infty} \frac{< h, v_i >^2}{\theta_i} := ||h||_{\mathbb{K}}^{2} < \infty\}$$

Here $K: \mathbb{H} \to \mathbb{H}$ is a positive definite, self-adjoint linear operator with spectral decomposition:

$$K = \sum_{i=1}^{\infty} \theta_i v_i \otimes v_i$$

where $\theta_i \geq 0$ and $v_i \in \mathbb{H}$ are the corresponding eigenvalues and eigenfunctions of $K$ respectively. This approach uses the norm $||\cdot||_{\mathbb{K}}$ to induce sparsity and smoothness, the degree of which is controlled using the tuning parameter $\lambda$. Note that for our study we followed the default for FLAME taking $K$ to be the Sobolev kernel.

We took $Y$ to be the growth curves (computed through the weight-for-length/height ratios in our main results) and $X$ a design matrix with each column corresponding to one of the 10,000 SNPs provided from screening. We standardized $X$ and centered $Y$ prior to applying FLAME. To tune the penalty for sparsity and smoothness, we split our observations into training (75%) and test (25%) sets and selected the $\lambda$ with minimum error on the test set. This procedure resulted in 24 SNPs (from 10,000) and their corresponding estimated effect curves.

We also used FLAME to assess the statistical robustness of SNP selection through a 20-fold sub-sampling scheme (selection was repeated 20 times, each time on 19/20 of the data). To compare selection results produced through similar amounts of induced sparsity and smoothness, we took $\lambda$ to be the parameter that originally produced the 24 SNPs in our PRS. The top 5 SNPs in terms of selection frequency and weight magnitude were taken to create FDA5 PRS. Due to the standardization step required for FLAME, final weights were adjusted by using the estimated effect curves produced when regressing the growth curves on the raw counts corresponding to our 24 and 5 selected SNPs.
